# Effects of deleterious mutations on the fixation of chromosomal inversions on autosomes and sex chromosomes

**DOI:** 10.1101/2025.10.10.681595

**Authors:** Denis Roze, Thomas Lenormand

## Abstract

Deleterious mutations have multiple effects on the fate of chromosomal inversions. In this article, we use individual-based simulations to estimate the fixation probabilities of neutral inversions under a constant input of deleterious mutations. As shown previously, we find that ‘lucky’ inversions carrying a lower-than-average mutation load are initially favored and tend to selectively spread. Our results also outline the importance of Muller’s ratchet caused by the absence of recombination in Y-linked and in rare autosomal inversions, reducing their fixation probabilities. Despite the fact that Y-linked inversions are more susceptible to Muller’s ratchet, they can more easily fix than autosomal inversions despite initially carrying mutations, when these mutations have sufficiently low selection or dominance coefficients (sheltering effect). Similarly, the sheltering of deleterious alleles can increase the fixation probability of inversions capturing a mating-type locus, particularly under intermediate rates of self-fertilization. Overall, our results confirm that when mating is random and for realistic parameter values, the presence of deleterious alleles reduces the average fixation probability of autosomal, sex-linked or mating type-linked inversions below that of a neutral mutation. Nevertheless, the occasional fixation of a lucky inversion may have important macroevolutionary consequences, potentially contributing to the evolution of recombination arrest on sex chromosomes.

## INTRODUCTION

Chromosomal inversions represent a common source of genetic polymorphism within populations and of fixed differences between species, and can play important roles in adaptation and speciation (Krimbas and Powell, 1992; Hoffmann and Rieseberg, 2008; Kirkpatrick, 2010; Wellenreuther and Bernatchez, 2018; Mérot et al., 2020). Inversions occur frequently (Ginner-Delgado et al., 2019; Porubsky et al., 2022) and may spread through different mechanisms (Kirkpatrick and Barton, 2006; Berdan et al., 2023). In a panmictic population, a newly arisen inversion should be present mostly in the heterozygous state (in heterokaryotypes). In this case, crossovers occurring in the region encompassed by the inversion may lead to large deletions or duplications that should strongly impair fitness, particularly in the case of pericentric inversions (that include the centromere; Sturtevant and Beadle, 1936; Hawley and Ganetzky, 2016). For this reason, inversions were previously thought to be underdominant (reducing the fitness of heterokaryotypes), requiring sufficiently strong genetic drift to increase in frequency (White, 1978; Lande, 1985). However, it was later shown that heterokaryotypes may not necessarily suffer from reduced fitness (e.g., Coyne et al., 1991, 1993), possibly due to the suppression of recombination within heterozygous inversions (Koury, 2023; Li et al., 2023). When an inversion captures a chromosomal segment that is fitter than the population average, it is selectively favored. This advantage may stem from a mutational effect caused by the inversion itself, or from a change in gene expression near one of its breakpoints (Wright and Schaeffer, 2022). Alternatively, it can result from the fact that the inversion has fortuitously captured a beneficial mutation or a beneficial combination of alleles at multiple loci, subsequently maintained in linkage within the inverted region. The last situation may involve epistatically interacting alleles (a “coadapted gene complex”; Dobzhansky, 1948, 1950; Charlesworth and Charlesworth, 1973), or a combination of locally beneficial alleles in an heterogeneous environment (Kirkpatrick and Barton, 2006; Charlesworth and Barton, 2018; Mackintosh et al., 2024). An inversion may also benefit from a fitness advantage if it carries fewer deleterious mutations than the population average. However, this advantage tends to erode over time due to the constant input of new deleterious mutations (Nei et al., 1967; Kimura and Ohta, 1970). Approximating this fitness erosion by a deterministic process, Connallon and Olito (2022) showed that, on average, the fixation probability of new inversions under a constant input of deleterious mutations stays close to the neutral expectation.

Inversions may play an important role in the evolution of sex chromosomes. Indeed, recombination arrest between the X and Y (or Z and W) chromosomes can be caused by inversions capturing the sex determining locus (e.g., Lahn and Page, 1999; McAllister, 2003; da Rosa et al., 2006). As in the autosomal case, such inversions may fix by drift (Ironside, 2010), or due to a selective advantage. In particular, an inversion that permanently links a male-beneficial allele to the male-determining allele (or a female-beneficial allele to the female-determining allele) benefits from a fitness advantage, corresponding to a particular case of the scenario of epistatically interacting mutations mentioned above. This corresponds to the classical theory to explain the early evolution of Y or W chromosomes, based on selection for suppressed recombination between sex chromosomes in order to permanently link sexually antagonistic loci to the sex determining locus (Rice, 1987, 1996; Charlesworth et al., 2005; Bachtrog, 2013), or more generally link the sex-determining locus with loci exhibiting sex differences in selection (Charlesworth and Charlesworth, 1980; Lenormand, 2003). More recently, several authors considered a scenario in which inversions fix on Y (or W) chromosomes because they carry a lower load of deleterious mutations than the population average (the “lucky inversion” scenario; Lenormand and Roze, 2022; Olito et al., 2024; Jay et al., 2022, 2024, 2025). As mentioned above, this benefit is only transient, since the marginal fitness of a mutation-free inversion tends to return to the population average as it reaches mutation - selection balance (Olito et al., 2024). Furthermore, an inversion fixed on the Y (or W) chromosome and capturing the sex determination locus never recombines (as it is only present in heterokaryotypes) and is thus prone to deleterious mutation accumulation by Muller’s ratchet (Muller, 1964; Felsenstein, 1974; Haigh, 1978). This should eventually favor the restoration of recombination (either by a re-inversion or by the evolution of a new sex determining locus elsewhere in the genome), unless the silencing of Y-linked genes and dosage compensation evolve sufficiently fast to stabilize recombination arrest (Lenormand and Roze, 2024).

Several authors proposed that inversions capturing the sex determining region of a Y or W chromosome may benefit from a selective advantage arising from their enforced heterozygosity (e.g., Branco et al., 2017; Ponnikas et al., 2018; Jay et al., 2022, 2024, 2025). This “sheltering” hypothesis proposes that recessive deleterious mutations present in such an inversion stay heterozygous, and therefore have less fitness effect than mutations present in a recombining region. This enforced heterozygosity of mutations would, in turn, facilitate the spread of inversions arising on Y (or W) chromosomes. In the context of sex chromosome evolution, the term “sheltering” initially appeared in articles considering the accumulation of deleterious mutations on an already non-recombining Y chromosome. In particular, in response to Muller’s (1914) verbal argument stating that the spread of fully recessive mutations on a Y chromosome should not be opposed by selection, Fisher (1935) showed that in infinite, randomly mating populations, the equilibrium frequency of a recessive lethal mutation on a Y chromosome is the same as on an autosome (due to the fact that the mutation also occurs on the X chromosome). Nei (1970) later showed that in small populations, the fixation probability of a fully recessive lethal mutation on a Y chromosome may approach the fixation probability of a neutral mutation (as the mutation is often absent from the population of X chromosomes, due to random drift). However, the fixation probability decreases rapidly when the mutation is not fully recessive and/or as population size increases. As discussed in Olito et al. (2024), several authors have mentioned the hypothesis that the sheltering of deleterious mutations may favor recombination arrest between sex chromosomes, referring to a previous model by Charlesworth and Wall (1999). This model showed that under partial sib-mating, linkage between the sex determining locus and a locus with heterozygote advantage is selectively favored. Indeed, under inbreeding, linkage to the sex determining locus increases heterozygosity, which is beneficial when heterozygotes have a higher fitness than homozygotes. Charlesworth and Wall (1999) conjectured that the same mechanism may favor recombination arrest when the higher fitness of heterozygotes is due to partially recessive deleterious mutations segregating at multiple loci, but did not further explore this scenario.

Jay et al. (2022) used a multilocus simulation approach to compare the rates of fixation of autosomal and Y-linked inversions, in the presence of recurrent deleterious mutations and under random mating. They observed higher rates of fixation of Y-linked inversions, and interpreted this result as an effect of the enforced heterozygosity of mutations present in Y-linked inversions (sheltering effect). Olito and Charlesworth (2023) and Charlesworth and Olito (2024) argued that the fact that Jay et al. (2022) show fixation rates of inversions that have survived for at least 20 generations (in particular, in their Figure 3c) gives the false impression that the fixation probability of Y-linked inversions is much higher than neutral, and may generate a bias increasing the fixation rate of Y-linked inversions relative to autosomal ones. Adding neutral controls (from simulations in which deleterious alleles are absent), Olito and Charlesworth (2023) showed that the fixation probability of Y-linked inversions is generally close to neutral or lower, except in restrictive conditions when the dominance coefficient of deleterious alleles is very low (a conclusion confirmed by analytical approximations derived by Olito and Charlesworth, 2023, showing that Y-linked inversions may then benefit from a net fitness advantage). These neutral controls also show that in some cases (in particular, when the fitness effect of mutations is very small), the increased fixation rate of Y-linked inversions relative to autosomal ones may be simply explained by their higher initial frequency (as neutral inversions display the same increased rate of fixation when they are Y-linked, see Figure 1 in Olito and Charlesworth, 2023). These criticisms ultimately led to the retraction of Jay et al. (2022). In a new version of their article, Jay et al. (2025) provided additional simulation results, confirming that the fixation probability of new inversions may be higher than neutral on Y chromosomes while being lower than neutral on autosomes when a substantial proportion of deleterious mutations are fully or almost fully recessive (*i*.*e*., about 30% of mutations have either *h* = 0 or *h* = 0.01; Figure 4 in Jay et al., 2025). However, the fact that Jay et al. performed their simulations over a fixed number of generations (10,000) complicates the interpretation of their results, as it generates a possible bias against the fixation of autosomal inversions relative to Y-linked ones, since fixation time should be longer in the case of autosomal inversions. Jay et al. (2025) also argued that the sheltering effect may increase the fixation probability of Y-linked inversions relative to autosomal ones even when these fixation probabilities are lower than neutral on average. In parallel, Olito et al. (2024) performed simulations under the simplifying assumption that the relative fitnesses of inversions follow deterministic trajectories until they reach mutation-selection balance (as in Connallon and Olito, 2022). They found similar fixation probabilities (relative to neutral) of new inversions on autosomes and on Y chromosomes, but did not consider the case of very recessive mutations (the dominance coefficient of deleterious alleles being set to either *h* = 0.25 or *h* = 0.1).

**Figure 1.**
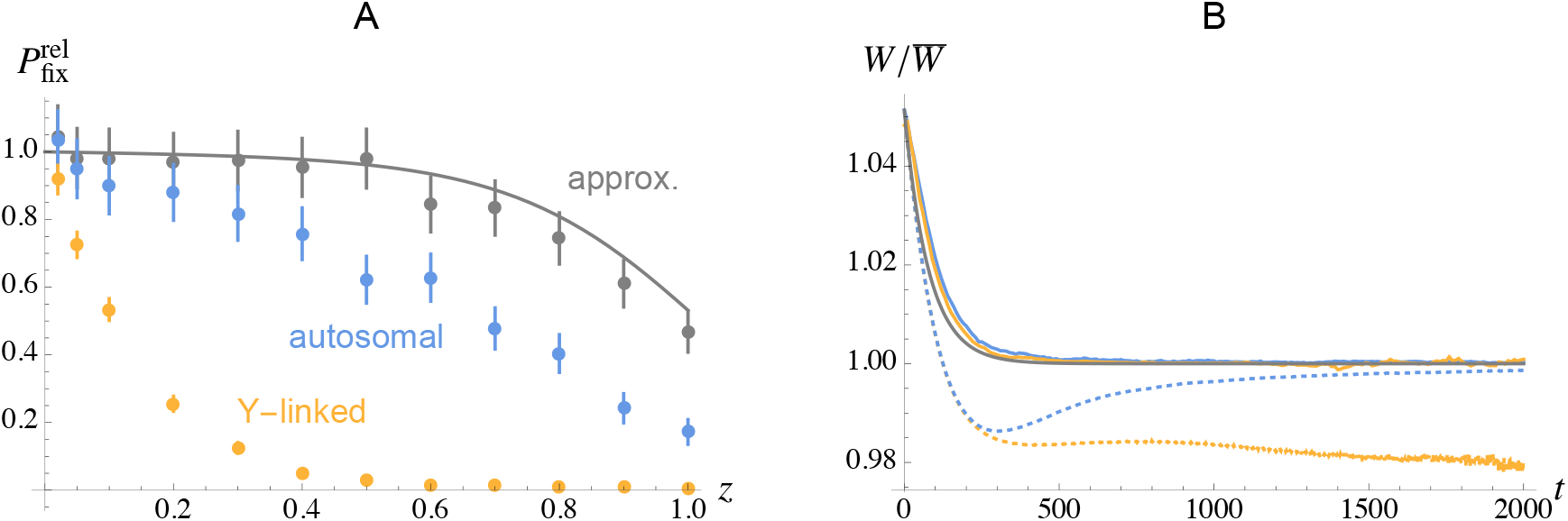
A. Fixation probability of an inversion of size *z* (measured as a fraction of the size of the chromosome) relative to the fixation probability of a neutral mutation. Grey curve: Kimura and Ohta’s (1970) approximation for autosomal inversions (obtained from their equations 8 and 9). Grey dots: stochastic simulations for autosomal inversions, assuming a deterministic change in frequency of their marginal fitness, as in Connallon and Olito (2022) and Olito et al. (2024). Blue dots: multilocus, individual-based simulations of autosomal inversions (see Methods). Orange dots: multilocus, individual-based simulations of Y-linked inversions (capturing the sex determining locus). Parameter values: *N* = 10,000, *U* = 0.1, *s* = 0.05, *h* = 0.25, *R* = 0.5. In this and the following figures, error bars on fixation probabilities correspond to support limits obtained by maximum likelihood (see Methods). B. Solid curves: mean relative marginal fitness of inversions as a function of time (in generations) in the multilocus simulations, averaged over the inversions of size *z* = 0.5 that reached fixation in the multilocus simulations: 311 autosomal inversions (blue), 64 Y-linked inversions (orange). Grey: deterministic prediction 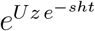, in the case of inversions that are initially mutation free (from Nei et al., 1967). Dotted curves: mean relative marginal fitness over time of inversions of size *z* = 0.5 that are initially mutation-free, including inversions that are eventually eliminated from the population. Averages are taken over ~ 10^7^ trajectories, the average at time *t* being measured over inversions that are still segregating at time *t* (~ 30, 000 autosomal inversions were still segregating at generation 2,000, but only ~ 500 Y-linked inversions).

These previous studies leave open a number of questions regarding the effect of deleterious mutations on the spread of inversions on autosomes and sex chromosomes. First, analytical treatments have provided important insights on the effect of deleterious alleles on the dynamics of inversions in large populations, but do not consider the effect of drift on changes in frequency of those alleles (Nei et al., 1967; Kimura and Ohta, 1970; Connallon and Olito, 2022; Olito et al., 2024). However, a rare autosomal inversion is mostly present in heterokaryotypes and therefore rarely recombines, generating interference among deleterious mutations present in the inversion, and possibly mutation accumulation through Muller’s ratchet. The consequences of interference should be far more important in the case of Y-linked inversions, as they never recombine. How does interference among mutations affect the fixation probability of inversions on autosomes and sex chromosomes? Second, is the sheltering advantage of Y-linked inversions only restricted to the case of fully or almost fully recessive deleterious alleles? How is it affected by the strength of selection against those alleles? Third, does the process described by Charlesworth and Wall (1999) also generate a selective advantage for inversions capturing a sex determining or mating type locus in an inbred population, in the presence of partially recessive deleterious mutations? In this article, we use individual based, multilocus simulations over a wide range of parameter values to answer these questions. Our results show that interference among mutations may significantly reduce the fixation probability of inversions, especially when they are Y-linked. However, this stronger effect of interference on Y-linked inversions may be compensated by the sheltering effect in regimes where inversions may reach fixation even when they initially carry deleterious mutations (which occurs when the effect of deleterious alleles is sufficiently weak). Finally, we show that partially recessive deleterious mutations can increase the fixation probability of inversions capturing a mating-type locus under intermediate selfing rates, but not under complete selfing.

## METHODS

### General framework

Our simulation programs (written in C++ and available from Zenodo, https://doi.org/10.5281/zenodo.17287166) represent a population of *N* diploid individuals with discrete generations. Each individual carries two copies of a linear chromosome, along which deleterious mutations occur at a rate *U* per chromosome per generation. All mutations have the same selection and dominance coefficients (*s* and *h*, respectively), although some simulations include two different types of mutations in varying proportions. Mutations have multiplicative effects on fitness (no epistasis), the fitness of an individual being given by:

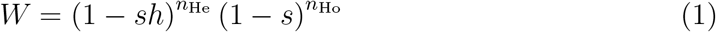

where *n*_He_ and *n*_Ho_ are the numbers of heterozygous and homozygous mutations present in its genome. Each generation, the number of new mutations on each chromosome is drawn from a Poisson distribution (with parameter *U*), the position of each new mutation along the chromosome being drawn from a uniform distribution (the number of sites at which mutations can occur is thus effectively infinite). The number of crossovers occurring at meiosis is drawn from a Poisson distribution with parameter *R* (chromosome map length), the position of each crossover being drawn from a uniform distribution (no interference). *R* is generally fixed to 0.5, corresponding to an average of one crossover per bivalent. However, different values of *R* were used for simulations performed under different values of the deleterious mutation rate *U*, in order to maintain a constant effective population size *N*_e_ in the absence of segregating inversion, taking background selection into account. Indeed, when *sh* ≪ *R* and *N*_e_ *sh* ≫ 1, equation 9 in Hudson and Kaplan (1995) leads to *N*_e_ ≈ *N e*^−2*U/R*^, showing that the effect of background selection depends on the *U/R* ratio. The population is initially mutation free, and is let to evolve during 2,000 generations to reach mutation-selection balance, before inversions are introduced. The number of preliminary generations was increased to 3,000 for the smallest value of *sh* considered (*sh* = 5 *×* 10^−4^).

### Autosomal case

Our autosomal model corresponds to the case of a standard Wright-Fisher population. After the preliminary generations, an inversion of size *z* (measured as a fraction of the total length of the chromosome, and fixed for each simulation) is introduced at a random location on a random chromosome. For this purpose, the position *x* of the left breakpoint of the inversion is drawn from a uniform distribution between 0 and 1 − *z*, the right breakpoint being at position *x* + *z*. The position of the inversion along the chromosome may affect its fixation probability only through potential effects of linked recombining regions, but we will see later that these effects are negligible in most cases. In heterozygous individuals for the inversion, recombination does not occur over the chromosome segment encompassed by the inversion, while the genetic map length outside the inversion remains equal to *R* (recombination is thus increased outside the inversion). Recombination occurs normally in individuals homozygous for the inversion. In some simulations, we included a rate *ρ* of double crossovers in heterokaryotypes. In this case, for simplicity the positions of both crossovers are drawn from a uniform distribution over the inversion length. The case *z* = 1 was considered as a theoretical limit: in this case, recombination is suppressed between inverted and non-inverted chromosomes over all of their length. The inversion is assumed to have no direct fitness effect, either in the heterozygous or homozygous state. The simulation continues until the inversion is eliminated from the population or reaches fixation; a new inversion is then introduced at random, and the whole process is repeated (generally 10^7^ times) to estimate fixation probabilities and times. Given that fixations correspond to binomial draws, support limits of the fixation probabilities estimates were computed as the values below and above the maximum likelihood estimate causing log-likelihood to drop by 1.92 units. The simulation program also records the marginal fitness of each inversion and the mean and variance in the number of mutations it carries every 10 generations after its birth and until fixation or elimination. Marginal fitness is measured by sampling 500 chromosomes carrying the inversion at random (with replacement), pairing them with randomly sampled chromosomes, and using equation 1 where *n*_He_ and *n*_Ho_ are now the numbers of heterozygous and homozygous mutations present in the genomic segment corresponding to the inversion. This marginal fitness is divided by the same quantity measured over randomly sampled individuals, to obtain the relative marginal fitness of inversions.

For most parameter values considered, deleterious alleles did not fix during the simulations; however, some fixations occurred for the smallest selection coefficients considered. In these cases, fixed mutations were removed from the population (every 200 generations, and every time an inversion reached fixation), in order to increase execution speed and improve our estimates of the average number of segregating deleterious alleles initially present in inversions that reach fixation.

### Inversion capturing a sex-determining locus

We assume here that sex is determined by a single locus present at the mid-point of the chromosome (in position 0.5), with a dominant male-determining allele (XY system), and consider inversions that capture the male-determining allele (all results also apply to the case of ZW sex determination and an inversion capturing the female-determining allele). The left breakpoint of an inversion of size *z* is drawn from a uniform distribution between 0.5 − *z* and 0.5 when *z* ≤ 0.5, and between 0 and 1 − *z* when *z >* 0.5, so that the inversion includes the sex-determining locus. Note that the inversion is only present in males and necessarily stays heterozygous, and therefore does not recombine. The program proceeds as in the autosomal case, except that when an inversion fixes (*i*.*e*., is present in all males), it is eliminated from all individuals with all the mutations it may contain, and the program performs 1,000 additional generations before introducing the next inversion (2,000 for the smallest value of *sh* considered), in order to restore a normal state of mutation-selection balance on a recombining chromosome (however, we observed that these additional generations have very little effect on the results).

### Inversion capturing a mating type locus

A third program considers the case of a mating type locus with two alleles (say + and −). Fusion is possible only between + and − gametes, so that all individuals are necessarily heterozygous at the mating type locus, which is located at the mid-point of the chromosome. The program proceeds as in the case of an inversion capturing a sex-determining allele, each new inversion capturing the + allele (and necessarily staying heterozygous). We also explore the effects of two forms of self-fertilization: “standard” self-fertilization, where each newly formed individual has a probability *σ* of resulting from the fusion of two gametes (carrying different mating type alleles) produced by two independent meioses from the same diploid parent, and automixis, where each new individual has a probability *σ* of being formed by two gametes produced by the same meiosis (and also carrying different mating type alleles).

### Effect of the recombining part of the chromosome

Different extensions of the programs are used to explore the effects of the recombining region of the chromosome on the spread of inversions. A first extension eliminates all mutations in the chromosomal region outside the inversion (this chromosomal region is not represented in this modified program), thereby eliminating any possible effect of the recombining region adjacent to the inversion. Comparisons with the results from the baseline program therefore allow us to test for a possible effect of deleterious alleles segregating in adjacent regions on the spread of inversions. In a second extension, deleterious mutations are still segregating over the whole chromosome, but the presence of an heterozygous inversion does not result in an elevated recombination rate in the remaining portion of the chromosome (the total genetic map length of the recombining region being *R* (1 − *z*) in heterokaryotypes, instead of *R* as in the baseline program). In this case, comparisons with the baseline program allow us to test to what extent the effect of an heterozygous inversion on recombination outside the inversion may affect its fixation probability.

## RESULTS

The figures shown in this article represent the fixation probability of inversions divided by the fixation probability of a neutral mutation (equal to its initial frequency). This relative fixation probability (denoted 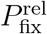) is thus given by 2*N P*_fix_, (*N/*2) *P*_fix_ and *N P*_fix_ in the case of autosomal inversions, inversions capturing the sex-determining locus on a Y-chromosome, and inversions capturing the + mating type allele, respectively, where *P*_fix_ is the absolute fixation probability. Because the rate of substitutions of inversions over time is given by the average number of new inversions per generation multiplied by their fixation probability (thus given by 2*Nµ*_A_ *P*_fix_, (*N/*2) *µ*_Y_ *P*_fix_ and *Nµ*_+_ *P*_fix_ in the same three cases, where *µ*_A_, *µ*_Y_ and *µ*_+_ are the rate of occurrence of the three types of inversion), the quantity 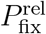 is thus equivalent to the rate of substitution, divided by the rate of inversion. Furthermore, given that the fixation probability of a neutral mutation is unaffected by any selection that might occur in the genetic background, 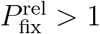 means that inversions are selectively favored on average (since they have a higher fixation probability than a neutral mutation), while 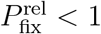 means that inversions are disfavored on average.

An inversion initially carrying a lower-than-average load of deleterious mutations is selectively favored, and tends to increase in frequency (Nei et al., 1967; Connallon and Olito, 2022; Lenormand and Roze, 2022; Olito et al., 2024; Jay et al., 2025). However, this initial advantage erodes over time as new mutations occur within the inversion. Previous studies obtained analytical and simulation results under the assumption that the change in the marginal fitness of inversions over time is deterministic (all copies of the inversion at a given generation having the same fitness). In particular, Nei et al. (1967) showed that, assuming that deleterious alleles are not too recessive and stay at low frequency, the relative fitness of an inversion of size *z* and initially free of deleterious mutation declines approximately as 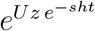, where *t* is time in generations (see also Connallon and Olito, 2022). This erosion of the initial fitness advantage of the inversion is due to the fact that the population of segregating copies of the inversion will eventually reach a state of mutation-selection balance. This result was incorporated into a diffusion model by Kimura and Ohta (1970) in order to derive fixation probabilities (which requires solving numerically an ordinary differential equation). More recently, Connallon and Olito (2022) obtained an approximation showing that, in the absence of any direct fitness effect of inversions and when the variance in fitness is generated by deleterious alleles at mutation-selection balance, the average fixation probability of new inversions should remain close to the fixation probability of neutral mutations. This result was confirmed by simulations following the stochastic fate of inversions in a finite population, but assuming a deterministic change in the marginal fitness of inversions over time (that is, without representing deleterious mutations explicitly; see also Olito et al., 2024).

Figure 1A compares the average fixation probability of inversions as a function of their size, obtained using Connallon and Olito’s simulation method for the case of autosomal inversions (using a similar program as in Connallon and Olito, 2022 and Olito et al., 2024, written in C++ and available from Zenodo), from Kimura and Ohta’s (1970) diffusion model, and from our multilocus simulation model in the case of autosomal and Y-linked inversions. We did not estimate the relative fixation probability of Y-linked inversions using Olito et al.’s semi-deterministic approximation, but they showed that it is very similar to the relative fixation probability of autosomal inversions obtained using the same method, for the parameter regime used in Figure 1 (Olito et al., 2024). Figure 1A shows that Kimura and Ohta’s method provides a reasonable match to the simulation results obtained using this approximation — Kimura and Ohta (1970) also proposed closed form approximations to their diffusion equation, but we found that their equation 11 (assuming *Uz* ≪ *sh*) does not hold for the parameter values considered here, while their equation 12 only holds for *z <* 0.25 (as can be deduced by the condition given by Kimura and Ohta), in which case the fixation probability stays very close to neutral. However, fixation probabilities are significantly lower in our multilocus simulations, particularly in the case of Y-linked inversions. Mutations segregating in the chromosomal region that always recombines (*i*.*e*., outside the inversion) do not contribute to this difference: indeed, Figures S1A and S2 show that the results obtained are very similar in the absence of deleterious mutations outside the inversion, for both the autosomal and Y-linked cases. Furthermore, Figure S1B shows that similar results are obtained when deleterious mutations occur a a finite number of sites regularly spaced along the chromosome (*L* = 1,000 sites with equal forward and back mutation rate *u* = *U/L*) instead of an infinite number of sites, showing that the difference is not caused by our assumption of an infinite number of sites at which deleterious mutations can occur.

Figure 1B shows the average relative marginal fitness over time of inversions of size *z* = 0.5 that reach fixation (solid curves on Figure 1B): the marginal fitness of fixed inversions is close to the deterministic prediction on average, but stays slightly higher, probably due to the fact that inversions that by chance accumulate mutations less rapidly tend to have higher fixation probabilities. As shown by Figure S3, this stochastic effect is relatively stronger for smaller inversions than for larger ones. However, while the marginal fitness of fixed inversions converges to the population average, inversions that do not reach fixation may decrease in fitness below the population average (even when initially favored), thus becoming disfavored. This is shown by the dotted curves on Figure 1B, representing the average fitness over time of inversions that are initially mutation-free, including inversions that are eventually eliminated from the population (averages are obtained from ~ 10^7^ trajectories, the average at time *t* being taken over all inversions that are still segregating after *t* generations). Figures 2A and 2C show 500 random trajectories of such initially mutation-free inversions, in the autosomal (2A) and Y-linked (2C) case. As can be seen on those figures, many trajectories show a rapid decline in fitness, the inversion becoming deleterious just before it is eliminated from the population. In the autosomal case, this effect is due to the fact that rare inversions stay heterozygous and therefore do not recombine, causing mutation accumulation by Muller’s ratchet (Muller, 1964; Felsenstein, 1974; Haigh, 1978): indeed, if the least-loaded class of inversions is lost by drift, it cannot be generated again by recombination when the inversion is only present in heterokaryotypes. Figures 2A and 2B show that the minimal number of deleterious mutations among all copies of the inversion present in the population does indeed increase along these declining trajectories, and that mutation accumulation occurs when inversions are below a threshold frequency (below ~ 0.1 in Figure 2B). Newly arisen inversions are particularly vulnerable to this process as they are initially in low frequency, causing a dip in average marginal fitness after ~ 300 generations (see blue dotted curve on Figure 1B). The effect of Muller’s ratchet is stronger in the case of Y-linked inversions since they never recombine, even when present at high frequency. Indeed, Figure 2D shows that mutation accumulation eventually occurs for all segregating inversions, even when they have reached high frequency. As a result, the mean fitness of segregating inversions is lower than in the autosomal case (compare blue and orange dotted curves on Figure 1B), explaining their lower relative fixation probabilities.

**Figure 2.**
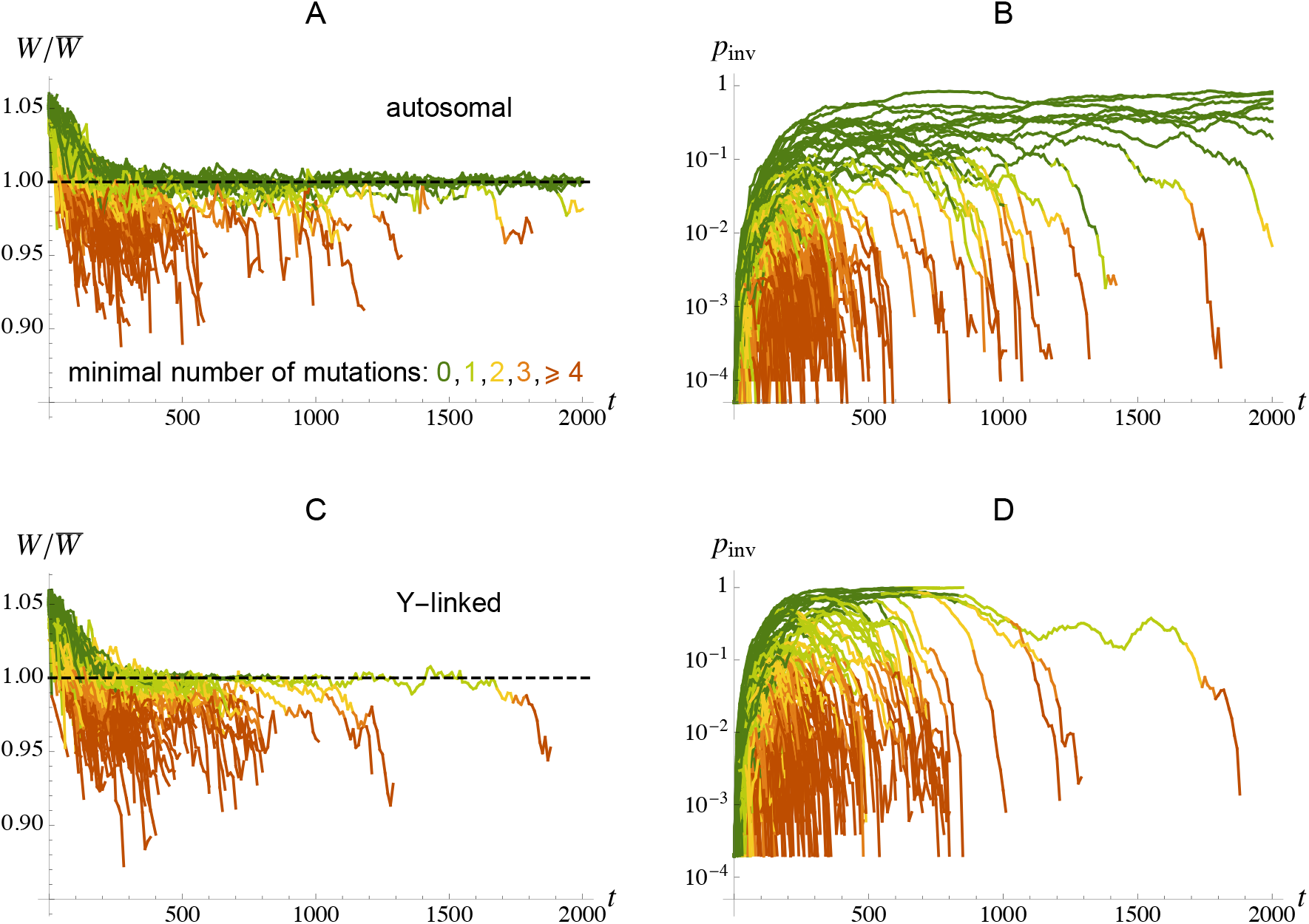
A, C: Relative marginal fitness over time of 500 inversions of size *z* = 0.5, initially mutation-free, in the autosomal (A) and Y-linked (C) cases (measured every 10 generations). Colors correspond to the minimal number of deleterious mutations per inversion, among all copies of the inversion present in the population (as indicated in A). B, D: Frequency of the inversion *p*_inv_ in the population over time, along the same trajectories as is A, C (and with the same color code).

Figure 3A shows the effect of population size *N* on the relative fixation probabilities of autosomal and Y-linked inversions of size *z* = 0.3, for the same parameter values as in Figure 1. Increasing population size increases the fixation time of inversions (which may decrease their fixation probability when their selective advantage declines over time), and decreases the effect of drift (which may increase the fixation probability of inversions benefitting from an initial advantage). Ignoring the effect of stochasticity on the dynamics of deleterious mutations (in the simulations or in the analysis) leads to the prediction that the second effect predominates, so that the relative fixation probability of inversions increases with population size (grey dots and curve in Figure 3A). The same effect of *N* is observed in our multilocus autosomal simulations (blue dots in Figure 3A). At the smallest population size considered (*N* = 200), Muller’s ratchet has no significant effect on the fixation probability of autosomal inversions (the blue and grey dots are superposed), as deleterious alleles do not have time to accumulate before inversions are fixed or lost from the population. However, Muller’s ratchet reduces the fixation probability of autosomal inversions at higher values of *N*, its effect being still visible at *N* = 10^5^. Indeed, beneficial inversions may accumulate deleterious alleles when they are still rare in the population, even when *N* is very large. In the case of Y-linked inversions, Muller’s ratchet causes a decrease in relative fixation probability as *N* increases up to ~ 2,000 (for the parameter values used in Figure 3A), presumably due to the increased fixation time of inversions, leaving more time for mutations to accumulate. The relative fixation probability then stays approximately constant between *N* = 2,000 to *N* = 10^5^, indicating that the effect of the longer fixation time of inversions may be compensated by the fact that Muller’s ratchet is slower when *N* is larger. Figure 3B shows the effect of introducing a rate *ρ* of double crossovers in heterokaryotypes (assuming for simplicity that the two crossovers occur at random locations within the inversion). As can be seen on the figure, double crossovers need to occur at a rate *ρ >* 0.01 (thus much higher than most estimates of the rate of double crossovers in heterokaryotypes, e.g., Guerrero et al., 2012) to significantly increase the fixation probability of Y-linked inversions.

**Figure 3.**
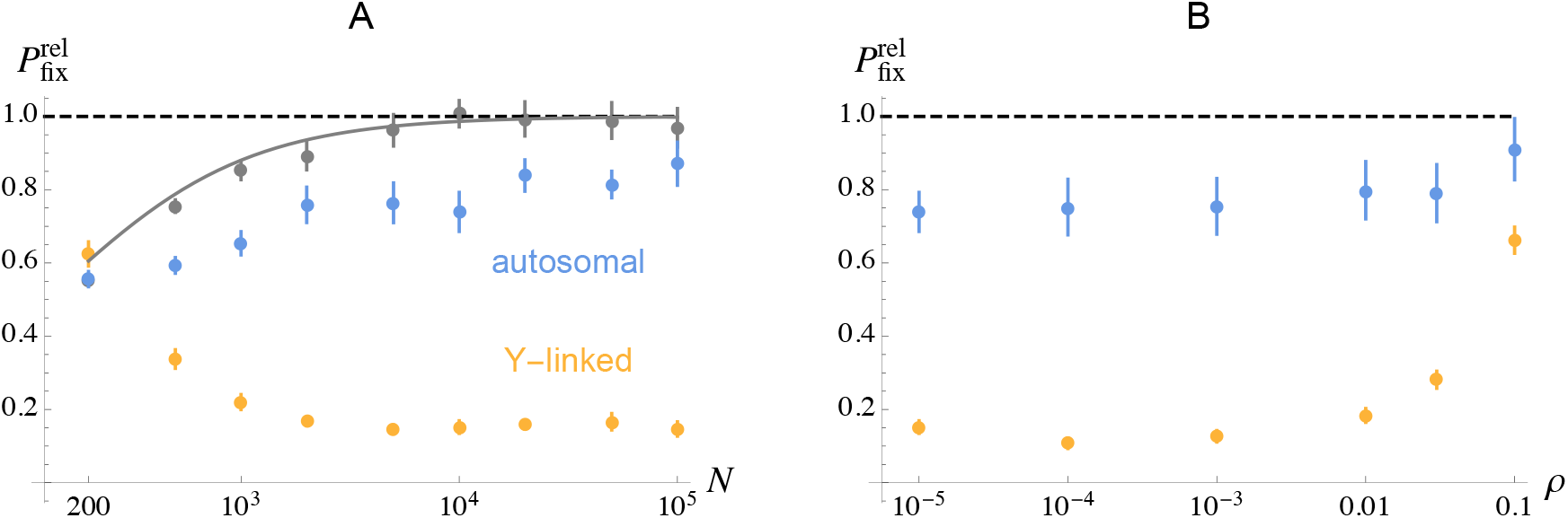
A. Fixation probability of an inversion of size *z* = 0.3 relative to the fixation probability of a neutral mutation, as a function of population size *N* (on a log scale). The color code is the same as in Figure 1A, and the curve also corresponds to Kimura and Ohta’s (1970) diffusion approximation for autosomal inversions (neglecting the effect of drift on the dynamics of deleterious alleles). In the multilocus simulations, mutations segregating outside the inversion are not included in order to reduce execution time. B. Fixation probability of an inversion of size *z* = 0.3 relative to the fixation probability of a neutral mutation, as a function of the rate of double crossovers *ρ* in heterokaryotypes (on a log scale). Parameter values are as in Figure 1, with *N* = 10^4^ in B.

Equations 8 and 9 in Kimura and Ohta (1970) predict that the fixation probability of an inversion of size *z* (relative to the fixation probability of a neutral mutation) should only depend on the two compound parameters *N*_e_ *Uz* (where *Uz* is the deleterious mutation rate within the inversion) and *N*_e_ *sh* (measuring the effective strength of selection against heterozygous deleterious alleles, relative to drift). The ratio of these two quantities affects the probability that a new inversion is mutation-free, which is approximately 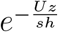 when selection against deleterious alleles is sufficiently strong relative to drift, so that the number of mutations per chromosome segment is approximately Poisson distributed. Additionally, *N*_e_ *Uz* affects the initial fitness advantage of a mutation-free inversion (given by *e*^*Uz*^), while *N*_e_ *sh* affects the rate of decay of this fitness advantage (given by *sh*, e.g., Nei et al., 1967). Numerical analysis of Kimura and Ohta’s diffusion equation indicates that the average fixation probability of inversions decreases as *N*_e_ *Uz* increases and as *N*_e_ *sh* decreases (results not shown). This indicates that decrease in the probability that a new inversion is mutation-free as *Uz* increases predominates over the effect of *Uz* on the initial fitness advantage of mutation-free inversions. Similarly, the increase in the probability that a new inversion is mutation-free as *sh* increases predominates over the effect of *sh* on the rate of fitness decay of mutation-free inversions. Alternatively, the results can be expressed in terms of the compound parameters *N*_e_ *sh* and *n*_mut_ = *Uz/* (*sh*), corresponding to the average number of mutations on a chromosome segment of size *z* at mutation-selection balance (when *sh* is not too small). In that case, Kimura and Ohta’s diffusion equation predicts that the average fixation probability of inversions should decrease as *n*_mut_ increases, as the probability that a an inversion is initially mutation free (given by 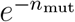) decreases. Under this parameterization and for a fixed *n*_mut_, increasing *N*_e_ *sh* has two contrasted effects: it increases the initial fitness of mutation free inversions (as *Uz* needs to increase with *sh* for *n*_mut_ = *Uz/sh* to remain constant), and it increases the rate of decay of this initial advantage. Numerical analysis of Kimura and Ohta’s diffusion equation indicates that the first effect predominates, so that the average fixation probability of inversions increases with *N*_e_ *sh* (for a fixed value of *n*_mut_).

The fact that fixation probabilities mostly depend on *n*_mut_ and *N*_e_ *sh* remains true in our multilocus simulations, in regimes when selection against deleterious alleles is sufficiently strong so that only the inversions that are initially mutation-free can reach fixation. This is shown in Figures 4A and 5: Figure 4A shows relative fixation probabilities (on a log scale) of autosomal and Y-linked inversions of different sizes *z* and for different deleterious mutation rates per chromosome *U* (for *Nsh* = 125), while Figure 5 shows results for different values of *s* and *h*. The *x*-axis of these figures (*n*_mut_) shows the average number of deleterious mutations on a chromosome segment of the size of the inversion (on a log scale): this quantity is measured in the simulations, but is very close to *Uz/* (*sh*) except when *h* is very low (Figures S4 and S5 show the same results as a function of inversion size *z*). Figures 5B, 5D and 5F show the average number of deleterious alleles initially present in inversions that reached fixation (*n*_init_). For the parameter values considered here, autosomal inversions that reach fixation are initially free of deleterious mutation in most cases, except in the case of relatively large inversions (roughly, *n*_mut_ *>* 10) when *h* = 0.5 and *s* = 0.005 or *s* = 0.001 (see red open circles on Figures 5D and 5F). In the regime where only the inversions that are initially mutation free may reach fixation, and for a given value of *Nsh*, Figures 4A and 5 show that fixation probabilities mostly depend on *n*_mut_, independently of *U*, *s* and *h*: *i*.*e*., open circles with different colors tend to fall along the same curves in Figures 4A and 5A, 5C, 5E (except when *h* = 0.5 and *n*_mut_ *>* 10). This is also true for Y-linked inversions, when *Nsh* = 125 (Figures 4A, 5A). By contrast, large Y-linked inversions can reach fixation even when they initially carry deleterious mutations, for *Nsh* = 25 and 5 and all values of *h* considered (Figures 5D – 5F). In this case, fixation probabilities do not depend only on *n*_mut_ and *Nsh* (*i*.*e*., filled circles do not fall along the same curves in Figures 5C and 5E), and may exceed the neutral fixation probability when *h* is very low (*h* = 0.004 or 0.02), indicating a net fitness advantage for Y-linked inversions. Figure S5 also shows the upper bound for the fixation probability of Y-linked inversions provided by equation 2 of Charlesworth and Olito (2024) when *h* = 0 and *N*_e_ *s* ≫ 1, which is significantly higher than our simulation results for *h* = 0.004. However, a better match is observed when this prediction is compared with simulations in which *h* = 0 and inversion size *z* is not too large, except in the case of recessive lethals (see Figure S6).

**Figure 4.**
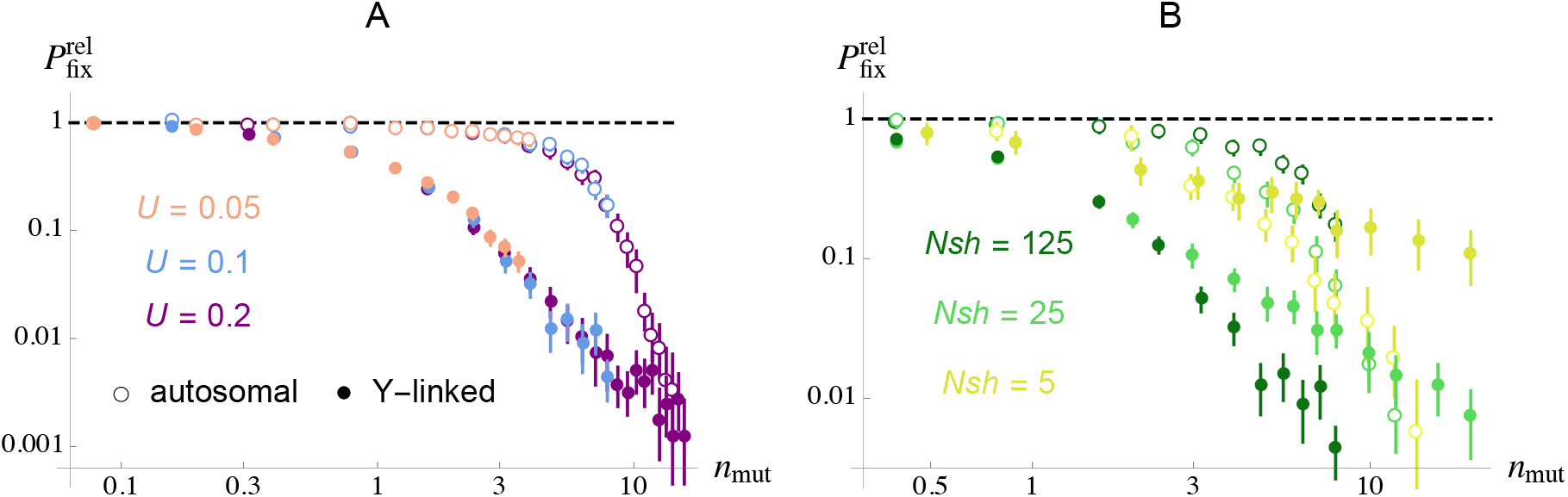
Fixation probability of autosomal and Y-linked inversions relative to the fixation probability of a neutral mutation (on a log scale), from multilocus, individual-based simulations. The x-axes show the average number of deleterious mutations in the population on a chromosome segment of the same size as the inversion (*n*_mut_, on a log scale). This number is measured in the simulations, but is generally very close to *Uz/* (*sh*). Empty and filled circles show results in the case of autosomal and Y-linked inversions, respectively. A. The different colors correspond to different rates of deleterious mutation per chromosome (*U* = 0.05, 1 or 2). For each color, the different points were obtained by performing simulations for different inversion sizes *z* (see Figure S4 for results as a function of *z*). B. The different colors correspond to different values of the strength of selection against deleterious alleles *s*, leading to different values of *Nsh*. Parameter values are *N* = 10^4^, *h* = 0.25, *s* = 0.05 (in A), *U* = 0.1 (in B). The chromosome map length *R* was set to 0.5 when *U* = 0.1 and to 0.25, 1 when *U* = 0.05, 0.2 (respectively) in order to maintain a constant *U/R* ratio and thus a constant *N*_e_ in the absence of segregating inversion (see Methods).

**Figure 5.**
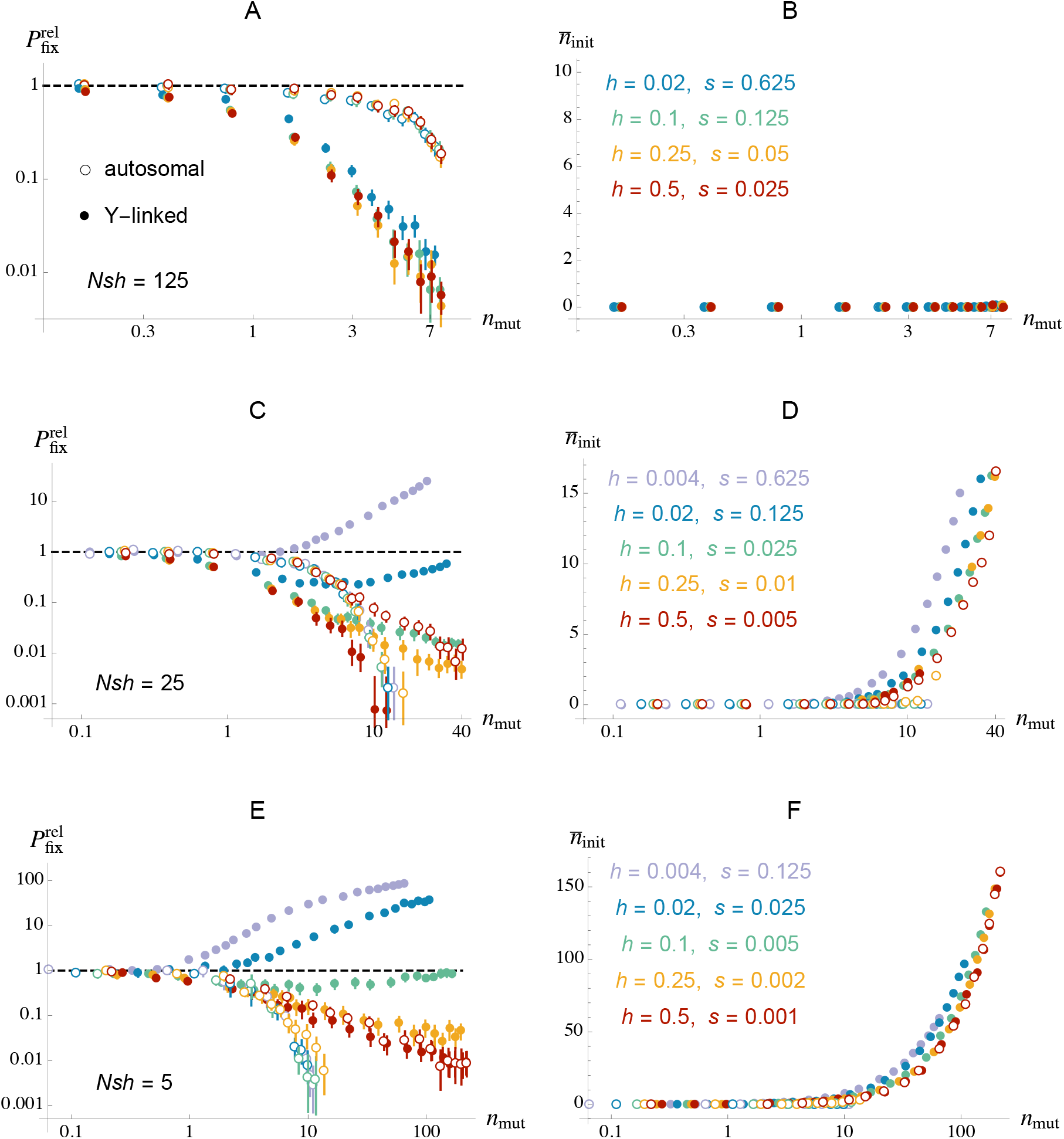
A, C, E: relative fixation probabilities of autosomal (empty circles) and Y-linked (filled circles) inversions as a function of *n*_mut_ ≈ *Uz/* (*sh*), the average number of deleterious mutations in the population on a chromosome segment of the same size as the inversion. As in Figure 4, for each color the different points were obtained by performing simulations for different inversion sizes *z* (see Figure S5 for results as a function of inversion size *z*). B, D, F: average number of deleterious alleles initially present in inversions that reached fixation (*n*_init_), over the same simulations as in A, C, E. Three values of *Nsh* were considered: *Nsh* = 125 (A, B), *Nsh* = 25 (C, D) and *Nsh* = 5 (E, F). In each case, different values of *h* (dominance coefficient of deleterious alleles) were considered, as shown in B, D, F. Other parameter values: *N* = 10^4^, *U* = 0.1, *R* = 0.5.

Figure 4B shows that in the case of autosomal inversions, and for a fixed *n*_mut_, the fixation probability of inversions tends to increase as *sh* increases, as predicted by Kimura and Ohta’s (1970) diffusion model. Furthermore, Figure S7 shows that *N* and *sh* mostly affect fixation probabilities through the *Nsh* product, as also predicted. In the case of Y-linked inversions, however, figures 4B and S7 shows that the effect of *Nsh* on fixation probabilities is opposite (*P*_fix_ increases as *Nsh* decreases, still for a fixed *n*_mut_). This may be due to the fact that the rate of fitness decline caused by Muller’s ratchet increases with the strength of selection against deleterious alleles (except when selection is very strong, Higgs and Woodcock, 1995; Gessler, 1995), and that Muller’s ratchet impacts Y-linked inversions more strongly than autosomal ones. Results on mean fixation times are shown in Figures S8 and S9: fixation times depend mostly on *n*_mut_ and *Nsh*, decreasing as *n*_mut_ increases (i.e., larger inversions tend to fix more rapidly) and as *Nsh* increases, for both autosomal and Y-linked inversions.

Overall, Figure 5 shows that while autosomal inversions have higher fixation probabilities (relative to neutral) than Y-linked inversions when *Nsh* is sufficiently large, Y-linked inversions have a higher relative fixation probability than autosomal ones when *Nsh* is smaller and *h <* 0.5, particularly in the case of large inversions and when *h* is small. In this second regime (weak selection against deleterious alleles, large inversions), virtually all new inversions carry deleterious mutations. Still, inversions carrying fewer mutations than the population average benefit from a fitness advantage, and tend to spread. In the autosomal case, this fitness advantage is reduced by the fact that, as the inversion increases in frequency, mutations that were initially present in the inversion necessarily become homozygous, and thus have a strong deleterious fitness effect. This generates a fitness cost for the inversion, explaining the drop in fixation probabilities observed in Figure 5 for *h <* 0.5 as the size of inversions (measured by *n*_mut_) increases. In the Y-linked case, however, inversions do not suffer from this cost of homozygosity and may fix even when they initially carry deleterious mutations (as shown by Figures 5D, 5F), even if their long-term marginal fitness is lower than the mean fitness of the equivalent portion of X chromosomes due to the presence of these initial mutations (Olito et al., 2024). Indeed, the marginal fitness of inversions changes slowly when *sh* is low (Nei et al., 1967), so that an inversion may fix before it has reached its equilibrium fitness. This corresponds to the sheltering effect described by Jay et al. (2025). As shown by Olito et al. (2024), mutations that are initially present in an inversion are also present in the population of X chromosomes (as these mutations were already segregating in the population when the inversion occurred) and may thus be homozygous, but the spread of the inversion tends to purge these mutations from X chromosomes (this can be seen on Figure S10A). Figure S11 shows that when the fixation of inversions that initially carry mutations is not taken into account, 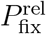 is similar or lower for Y-linked inversions than for autosomal ones (confirming that the higher 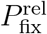 of Y-linked inversions observed in Figure 5 is due to the sheltering effect), except when *h* = 0.004 (in which case 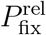 is slightly higher for Y-linked inversions). This modest increase in the relative fixation probability of mutation-free, Y-linked inversions (relative to autosomal ones) when *h* = 0.004 may stem from the enforced heterozygosity of new mutations arising in Y-linked inversions after their birth, in regimes where a substantial component of the mutation load of autosomes is caused by homozygous mutations (that is, when *h* is sufficiently close to zero, as shown by Figure S10B). As for the sheltering of mutations initially present in the inversion, this benefit is only transient, as enforced heterozygosity increases the equilibrium mutation load caused by nearly recessive mutations (Chasnov, 2000). Furthermore, Figure S12 shows that the net advantage of Y-linked inversions 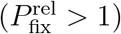 observed when deleterious alleles are very recessive is maximized at an intermediate value of population size *N*, and turns to an average disadvantage above a threshold value of *N*. This may be understood from the fact that the fixation time of inversions increases with *N*, leaving more time for mutations to accumulate.

Our infinite sites model of deleterious mutations may be seen as a best case scenario for the effect of sheltering, since a mutation present in a Y-linked inversion and that has been purged from the population of X chromosomes cannot re-occur on the X. In a model including recurrent deleterious mutations at a finite number of sites, a mutation present in a Y-linked inversion could become homozygous due to recurrent mutations on the X, thus reducing the fitness of the inversion (Fisher, 1935; Nei, 1970). Figure S13 shows that the sheltering effect observed for *h* = 0.004 and *s* = 0.625 is indeed lowered when mutations occur at *L* = 1,000 loci spread along the chromosome (leading to a per locus mutation rate of *u* = 10^−4^), while results obtained for *L* = 10,000 (and *u* = 10^−5^) are similar to those obtained under an infinite number of sites. The effect of recurrent mutations likely depends on the *Nu* product, becoming negligible when *Nu* is sufficient small (here *Nu* = 1 for *L* = 1,000, while *Nu* = 0.1 for *L* = 10,000). By contrast, *L* has no noticeable effect in regimes where sheltering does not occur (Figures S13A – S13C). In this case, the number of selected sites within an inversion should only have an effect through the overall deleterious mutation rate in the inversion, the effect of the per-locus mutation rate becoming more important in regimes where sheltering may occur.

Figure 6 shows relative fixation probabilities of autosomal and Y-linked inversions of size *z* = 0.1 and *z* = 0.2, when two types of deleterious mutations co-occur: mutations with *s* = 0.01, *h* = 0.25, coexisting either with very recessive mutations but with the same heterozygous effect (*s* = 0.625, *h* = 0.004, top figures), or with more weakly selected mutations but with the same dominance coefficient (*s* = 0.002, *h* = 0.25, bottom figures). While autosomal inversions of both sizes have a higher relative fixation probability than Y-linked inversions when *s* = 0.01 and *h* = 0.25 for all mutations (left-most points in Figure 6), the relative probability of fixation of Y-linked inversions becomes higher than autosomal inversions when the frequency of very recessive mutations (top figures) or weakly selected mutations (bottom figures) is sufficiently high, the threshold being reached more rapidly in the case of larger inversions (compare left and right figures). However, the average fixation probability of Y-linked inversions remains lower than neutral in most cases, except when all mutations are highly recessive (top figures with *p*_rec_ = 1).

**Figure 6.**
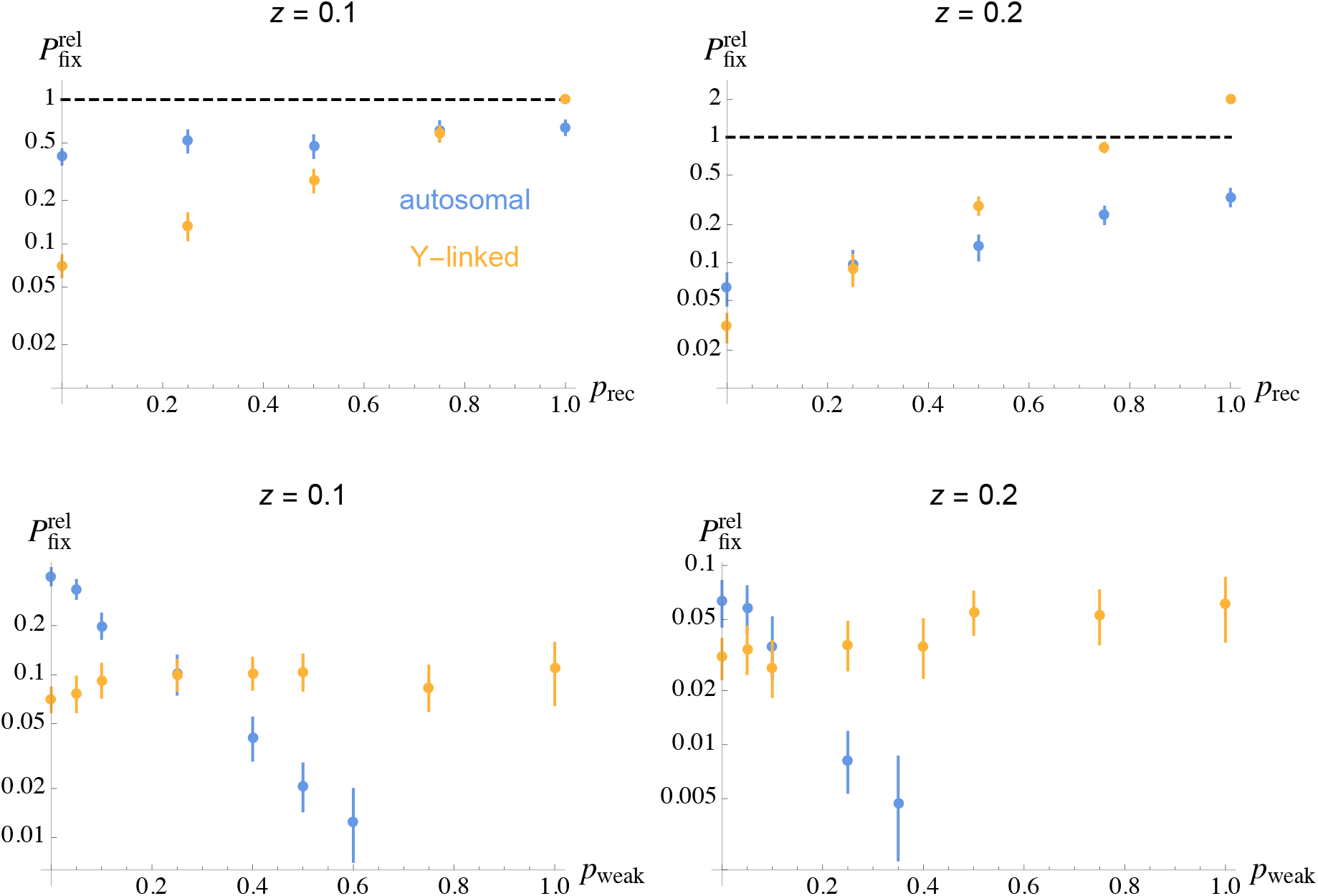
Relative fixation probabilities of autosomal (blue) and Y-linked (orange) inversions of size *z* = 0.1 (left) and *z* = 0.2 (right), when two types of deleterious mutations co-occur. In the top figures, a proportion 1 − *p*_rec_ of new mutations have *s* = 0.01 and *h* = 0.25, while a proportion *p*_rec_ have *s* = 0.625 and *h* = 0.004 (both types of mutations have the same heterozygous effect *sh* = 0.0025, but the second type is more recessive). In the bottom figures, a proportion 1 − *p*_weak_ of new mutations have *s* = 0.01 and *h* = 0.25, while a proportion *p*_weak_ have *s* = 0.002 and *h* = 0.25 (both types of mutations have the same dominance coefficient, but the second type is more weakly selected). Other parameter values are *N* = 10^4^, *U* = 0.1, *R* = 0.5.

In the absence of inbreeding, results on the fixation probability of inversions capturing a mating-type locus are qualitatively and quantitatively similar to the results obtained in the case of Y-linked inversions (compare Figure S14 and Figures 4A, 5A and 5C): indeed, both types of inversions are maintained permanently heterozygous, leading to similar advantages and disadvantages. As in the Y-linked case, the probability of fixation of an inversion capturing a mating-type locus is little affected by deleterious alleles segregating outside the inversion (Figure S15). However, in the presence of partial selfing or automixis, and when deleterious alleles are partially recessive (*h <* 0.5), inversions capturing a mating-type locus can become advantageous over a wider range of parameters than under random mating (Figure 7). This advantage becomes particularly strong when mutations are very recessive, for the smallest value of *Nsh* considered (*Nsh* = 25, lower panel of Figure 7). In the case of additive mutations (*h* = 0.5), however, the fixation probability of inversions capturing the mating-type locus decreases as the rate of inbreeding increases, while the effect of the selfing rate *σ* may be non-monotonic when *h <* 0.5. As shown by Figure S16, for *Nsh* = 125 fixation probabilities are lower at *σ* = 0.9 than at *σ* = 0.5 (where *σ* is the selfing rate) for all values of *h* considered, while for *Nsh* = 25 fixation probabilities are higher at *σ* = 0.9 than at *σ* = 0.5 for *h* = 0.1 and 0.25, but equivalent or lower for *h* = 0.004, 0.02 and 0.5. Additional simulations were performed under complete selfing (*σ* = 1). For *Nsh* = 125, mutations accumulated rapidly when *h* = 0.5, while mutation-selection balance was reached for lower values of *h* (and higher values of *s*); however, no fixation of inversion was observed over the 10^7^ trials performed for each parameter set, except for *h* = 0.25 and *z* ≤ 0.05 (leading to estimates of 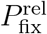 of 0.19 and 0.021 for *z* = 0.02 and *z* = 0.05, respectively). For *Nsh* = 25, mutations accumulated rapidly for all values of *h* except *h* = 0.004 and 0.02. In these cases, fixation probabilities were much lower than at *σ* = 0.9, and inversions of intermediate sizes did not fix, except under automixis with *h* = 0.02 (see Figure S17).

**Figure 7.**
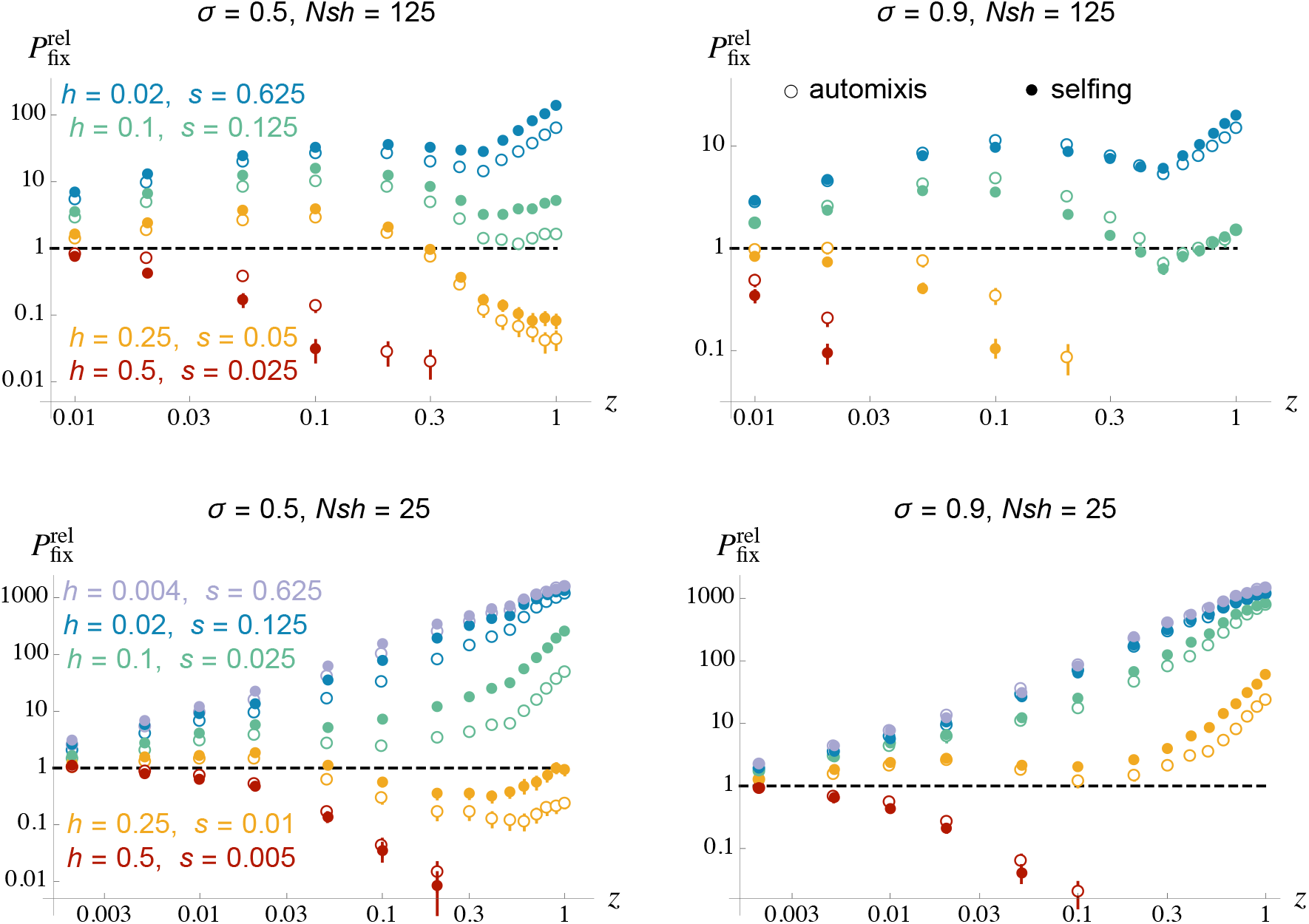
Relative fixation probabilities of inversions capturing a mating-type locus as a function of their size *z*, for different values of the rate of selfing or automixis *σ*, and different selection and dominance coefficients of deleterious alleles. Left: *σ* = 0.5; right: *σ* = 0.9. Filled circles correspond to partial selfing, open circles to partial automixis. Other parameter values are *N* = 10^4^, *U* = 0.1, *R* = 0.5.

These results show that inbreeding has contrasted effects on the spread of inversions capturing a mating-type locus. The increase in fixation probabilities observed under intermediate selfing rates and *h <* 0.5 is probably due to the masking of deleterious alleles either initially present or arising within spreading inversions. Indeed, recombination and inbreeding tend to generate homozygosity (so that homozygous mutations make a substantial contribution to the mutation load even when *h* is not very small), while recombination arrest maintains heterozygosity around the mating-type locus. However, homozygosity increases the efficiency of selection against deleterious mutations (for this reason, the equilibrium mutation load is lower under selfing than under outcrossing, e.g., Charlesworth et al., 1990). This reduction in load decreases the initial advantage of mutation-free inversions. Furthermore, an inversion capturing the mating-type locus is expected to be less well purged than the equivalent chromosome segment within individuals that do not carry the inversion, and its marginal fitness will thus eventually reach a lower value than the marginal fitness of recombining segments, even when the inversion is initially mutation-free. This is illustrated by Figure S18, showing that the initial fitness advantage of lucky inversions capturing the mating-type locus vanishes more rapidly under complete selfing (*σ* = 1) than under intermediate selfing (*σ* = 0.5), explaining the low fixation probabilities of such inversions under complete selfing. The effect of this reduction in purging of deleterious alleles (which disfavors inversions) is expected to become more important as *h* increases (since the masking advantage is stronger when *h* is lower), and as *s* increases (since the equilibrium mutation load is reached more rapidly).

Figure 8 shows fixation probabilities of inversions capturing a mating-type locus when *Nsh* = 25 and under intermediate selfing (*σ* = 0.5), for different proportions of strongly recessive mutations (*h* = 0.004). Average fixation probabilities are significantly increased when 5% or 10% of the mutations are strongly recessive, but generally stay of a similar order of magnitude as neutral mutations, on average. Figures S19 and S20 explore the effects of deleterious mutations segregating in adjacent, recombining chromosomal regions on the fixation probability of inversions under partial selfing. Contrarily to the results observed in the absence of selfing (Figure S15), deleterious mutations segregating in linked recombining regions tend to increase the fixation probability of inversions, when *h <* 0.5 (compare orange and grey dots in Figures S19 – S20). Furthermore, the effect of linked regions becomes stronger when recombination in these regions is decreased in heterokaryotypes (compare orange and green dots in Figures S19 – S20). A possible explanation for these results is that successful inversions tend to be initially associated with relatively fit linked regions, that can remain associated to the inversions during several generations due to selfing (particularly when recombination is low in these regions), and may also generate a masking advantage in the presence of partially recessive deleterious mutations.

**Figure 8.**
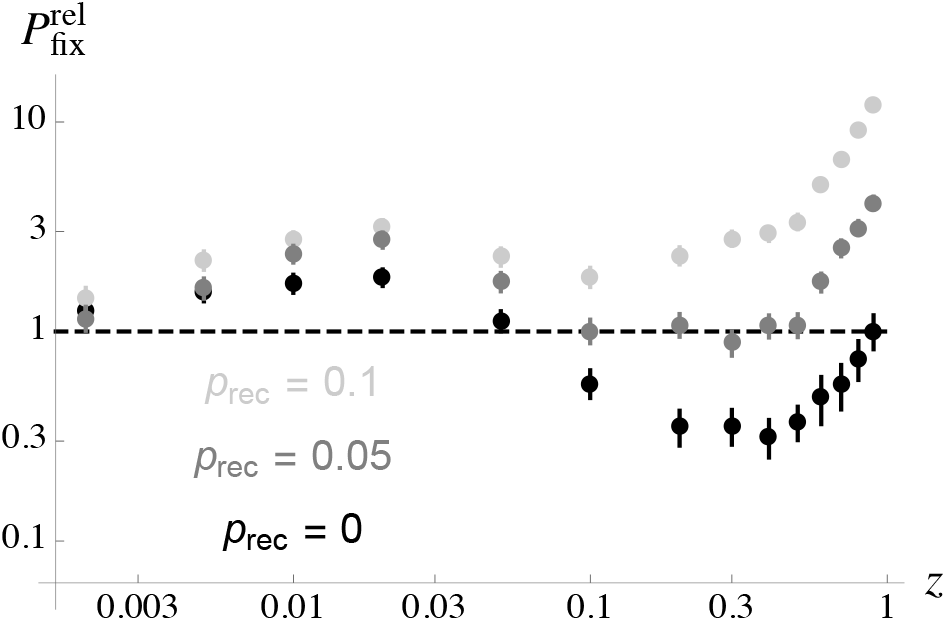
Relative fixation probabilities of inversions capturing a mating-type locus as a function of their size *z*, under partial selfing (*σ* = 0.5). Two types of deleterious mutations co-occur in the simulations: a proportion 1 − *p*_rec_ of new mutations have *s* = 0.01 and *h* = 0.25, while a proportion *p*_rec_ have *s* = 0.625 and *h* = 0.004 (both types of mutations have the same heterozygous effect *sh* = 0.0025, but the second type is more recessive). Other parameter values are *N* = 10^4^, *U* = 0.1, *R* = 0.5.

While regular self-fertilization (fusion between gametes from different meioses) and automixis (fusion between gametes from the same meiosis) have similar qualitative effects on fixation probabilities of inversions capturing the mating-type locus, quantitative differences can be observed on Figures 7 and S17. These differences must be due to the fact that, on the chromosome carrying the mating-type locus, homozygosity is lower under automixis than under regular selfing. This is most easily seen in the case of a focal locus which is tightly linked to the mating-type locus (but remains true in the case of more distant loci). Starting from an heterozygous parent at the focal locus, the probability that an offspring is homozygous at this locus is approximately 2*r* under regular selfing (where *r* is the recombination rate between the mating-type and focal locus, assumed small), corresponding to the probability that one of the fusing gametes is a recombinant; however, it is only *r* under automixis (the probability that a crossover occurs between the two homologs is approximately 2*r*, but only half of the zygotes possibly formed will be homozygous). This lower homozygosity (in the absence of inversion) reduces the masking advantage of inversions capturing the mating-type locus, but also reduces their disadvantage in terms of less efficient purging, explaining why fixation probabilities may be either higher or lower under automixis than under regular selfing depending on which of these two effects is strongest.

## DISCUSSION

Several recent models have quantified the fixation probabilities of inversions occurring in different parts of the genome, under various scenarios (Connallon and Olito, 2022; Olito et al., 2024; Jay et al., 2022, 2025). In general, fixation probabilities inform us about rates of evolutionary change over time (that depend on the rate of occurrence of new variants and their fixation probability), on the effective strength of selection (relative to drift) and overall direction of evolution in finite populations (e.g., Bulmer, 1991; Rousset, 2004; Lehmann and Rousset, 2009). In the context of the evolution of recombination, this approach has been used to quantify the effective strength of selection on mutations affecting the genetic map length of chromosomes, by comparing their fixation probability to the fixation probability of neutral mutations (e.g., Keightley and Otto, 2006; Hartfield et al., 2010). Regarding the evolution of Y (or W) chromosome map length, several authors have considered the fixation probability of inversions arising on sex chromosomes — typically on the Y, but inversions on the X can also lead to Y recombination suppression (Olito et al., 2024; Jay et al., 2025; Flintham and Mullon, 2025). This approach can be used (1) to characterize whether inversions can be selectively favored (by comparing their fixation probability to that of a neutral variant) and (2) to dissect the processes and causes influencing their fate (by comparing their fixation probability to that of inversions occurring in different genomic regions, notably on autosomes). However, it also presents several limitations that are useful to mention at the outset. First, average fixation probabilities indicate whether inversions are on average selectively favored or not, which is only partial information. Different inversions may have widely different fixation probabilities (depending in particular on their initial load of deleterious alleles) and their distribution is certainly important, especially if few of them can turn out to be selectively favored while most are not. Similarly, the average fixation probability of “mutations” occurring in the genome is likely to be lower than neutral, since most mutations with phenotypic effects are deleterious. Yet, it should not obscure the fact that a handful of beneficial mutations play a major role in adaptation. In the context of the fixation of Y-linked inversions, even if their average probability of fixation is (much) lower than neutral, many inversions may nevertheless fix on a sufficiently large time scale, particularly if inversions occur frequently (as suggested by recent data, e.g., Ginner-Delgado et al., 2019; Porubsky et al., 2022). For instance, in a regime where sheltering is not operating (*h* = 0.25, *Ns* large for most mutations), many inversions can fix (see Figures 2 and S2-S4 in Lenormand and Roze, 2022, Figure 3 in Lenormand and Roze, 2024). Given the rate of accumulation of non-recombining strata on sex chromosomes (for example, four strata on the mammalian Y chromosome in 180 millions years of evolution; Hughes and Page, 2015), invoking a process leading to a higher than neutral average fixation rate of Y-linked inversions may not be necessary — unless the restoration of recombination by re-inversions is so frequent that only a small proportion of fixed inversions may evolve as stabilized strata (Lenormand and Roze, 2024). Second, the fixation probabilities of mutations restoring recombination on sex chromosomes is usually not considered in comparison. Yet, they need to be accounted for to understand the long-term evolution of Y chromosome map length. We will return to these two issues through the following discussion focusing on the effects of deleterious mutations on the spread of chromosomal inversions.

### How do deleterious mutations affect the dynamics of chromosomal inversions?

Deleterious alleles have two main effects on the fate of inversions. As described by Nei et al. (1967), a first effect is that they generate a variance in fitness between copies of the same chromosome segment present in different individuals. Therefore, a new inversion may be favored or disfavored depending on its load of deleterious alleles relative to the population average: lucky inversions carrying a lower load than the population average tend to spread. However, this initial advantage erodes over time, as shown by deterministic models and by stochastic multilocus simulations (Nei et al., 1967; Connallon and Olito, 2022; Lenormand and Roze, 2022; Olito et al., 2024; Jay et al., 2025). Indeed, autosomal and Y-linked inversions that are initially free of deleterious mutation tend to deterministically return to the population average load, while inversions that initially carry one or several deleterious alleles will eventually become disfavored (Olito et al., 2024). However, inversions initially carrying mutations can still fix in parameter regimes allowing fixation to occur before the inversion becomes selectively disadvantageous. In our simulations, and in the case of autosomal inversions, this occurred mostly when mutations had weak and additive fitness effects (*s* = 0.001 or 0.005, *h* = 0.5). However, it occurred much more frequently in the case of Y-linked inversions due to the sheltering effect: mutations that are initially present in an inversion are mostly expressed in the heterozygous state even when the inversion is frequent, thereby causing a smaller disadvantage than in the autosomal case. Note that this effect does not provide a fitness advantage for Y-linked inversions (their advantage stems from their lower than average initial load), but reduces the disadvantage of initially carrying deleterious alleles. The values of *Nsh* at which we observed this effect (*Nsh* = 5 and 25) are lower than those considered by Olito et al. (2024) — for example, *Nsh* = 250 and 2,500 in their Figure 2 — explaining why the fixation of initially loaded inversions did not occur in their simulations. The second effect of deleterious mutations on the fate of chromosomal inversions is caused by Muller’s ratchet (Muller, 1964; Haigh, 1978): in a finite population and in the absence of recombination, deleterious mutations tend to accumulate over time, causing a steadily decline in fitness. In the case of autosomal inversions, this may cause a decrease in fitness of rare inversions (that are mostly present in heterokaryotypes and therefore do not recombine) and reduce their fixation probability. However, Muller’s ratchet may have a much stronger effect in the case of Y-linked inversions, as it may operate even in frequent or fixed inversions (Charlesworth, 1978). It is important to note here that our model focuses on inversions that would otherwise be neutral if no deleterious allele is segregating, and that the reduction in fixation probability caused by Muller’s ratchet may be less important in the case of inversions benefitting from an inherent fitness advantage (for example, inversions capturing a beneficial mutation or a beneficial combination of alleles). Indeed, such inversions are expected to rise in frequency more rapidly, reducing the risk of mutation accumulation while they are still rare. This could be further explored by extending our simulation framework to the case of inherently beneficial inversions.

### Do deleterious mutations help or hinder the fixation of inversions on autosomes and sex chromosomes, on average?

This can be deduced by the average fixation probability of inversions in the presence of deleterious mutations, compared to the fixation probability of a neutral mutation (as the latter corresponds to the fixation probability of a neutral inversion in the absence of deleterious mutations). As in previous studies (Connallon and Olito, 2022; Olito et al., 2024), we found that the fixation probability of small inversions is close to neutral, while the average fixation probability of larger autosomal and Y-linked inversions is generally lower than neutral under random mating, showing that the average effect of deleterious mutations is to hinder the fixation of inversions. Indeed, lucky inversions arising on a particularly good genetic background have higher than neutral fixation probabilities, but this effect is compensated by the fact that the probability that a new inversion is initially mutation-free decreases as the inversion’s size increases. Furthermore, Muller’s ratchet tends to reduce the fixation probability of inversions (in particular when they are Y-linked). However, it is important to note that although average fixation probabilities are nearly neutral or less than neutral, the rare inversions that manage to fix are selectively favored, and the process is therefore not neutral. For example, for *N* = 10^4^, *s* = 0.01, *h* = 0.25 and *U* = 0.1, an autosomal inversion spanning a fifth of the size of the chromosome (*z* = 0.2) has an average fixation probability of *P*_fix_ ≈ 3.2 *×* 10^−6^, which is much lower than the fixation probability of a neutral mutation (5 *×* 10^−5^). However, given that only the inversions that are initially mutation-free can fix for these parameter values (as shown by the simulations), and that the proportion of new inversions that are mutation-free is approximately *π*_0_ = *e*^−*Uz/*(*sh*)^ ≈ 3.35 *×* 10^−4^, one finds that the fixation probability of mutation-free inversions is *P*_fix_*/π*_0_ ≈ 9.5 *×* 10^−3^, which is about 190 times the fixation probability of a neutral mutation (while their mean fixation time is ≈ 0.074 times the mean fixation time of neutral mutations). Although these events are rare, they lead to the fixation of a whole chromosome segment and may thus have a significant effect on neutral diversity patterns in species in which inversions occur frequently. Quantifying the possible strength of this effect would represent an interesting extension of this work. In the case of Y-linked inversions, we found that the average fixation probability of inversions may be higher than neutral when mutations are almost fully recessive (*h* = 0.004, *h* = 0.02) in regimes where inversions may fix despite initially carrying mutations, in agreement with the results of Jay et al. (2025). This average advantage of inversions is due to the fact that the proportion of inversions that can maintain their selective advantage until fixation is enlarged in these regimes, since inversions may fix despite initially carrying mutations. Although the average fixation probabilities are less than neutral in all other cases investigated, this should not obscure the fact that most fixed inversions benefit from a selective advantage, as in the autosomal case discussed above.

### Does linkage to the sex-determining region (SDR) of Y or W chromosomes increase or decrease the mean disadvantage of chromosomal inversions caused by deleterious mutations?

This question is probably the main source of disagreement between Olito et al. (2024) and Jay et al. (2025), as Olito et al. (2024) predict that the fixation probability of autosomal and Y-linked inversions (relative to neutral) should be similar, while Jay et al. (2025) predict that linkage to the SDR increases the fixation probability of inversions. The average relative fixation probability of autosomal and Y-linked inversions can be compared in order to assess whether they tend to be more or less advantaged or disadvantaged depending on their genomic localization. Our results show that these average relative fixation probabilities generally differ, and that the sign of this difference depends on parameter values. Indeed, and as discussed above, linkage to the SDR has two contrasted effects: it may generate a sheltering advantage, but it also increases the effect of Muller’s ratchet. The first effect occurs in regimes where Y-linked inversions may fix despite initially carrying mutations, that is, when mutations are weakly selected and partially recessive. By contrast, the effect of Muller’s ratchet is expected to become stronger as selection against deleterious alleles increases (except in the case of very strong selection; e.g., Higgs and Woodcock, 1995; Gessler, 1995). As a result, we found that linkage to the SDR hinders the fixation of inversions (due to Muller’s ratchet) when *Nsh* is sufficiently large (*Nsh* = 125 in Figure 5A), while linkage to the SDR may help the fixation of chromosomal inversions (through the sheltering effect) under lower values of *Nsh*, especially in the case of large inversions and for low values of *h*.

Whether linkage to the SDR should help or hinder the spread of chromosomal inversions therefore depends on the distribution of fitness effects (DFE) and dominance coefficients of deleterious mutations, as well as on population size. Different methods have been used to estimate DFEs, leading to different estimates of the proportion of nearly neutral mutations (*N*_e_ *s* ~ 1): methods based on genetic polymorphism data typically infer higher proportions of nearly neutral mutations than methods based on mutation accumulation lines or on direct measurements of the fitness effect of induced mutations (e.g., Halligan and Keightley, 2009; Bataillon and Bailey, 2014; Charlesworth, 2015; Kim et al., 2017). Our simulations indicate that in a population of size 10^4^, with a deleterious mutation rate of 0.1 per chromosome and for *h* = 0.25, only large inversions have a higher relative probability of fixation on Y chromosomes than on autosomes when *s* = 0.01 (*Ns* = 100), while most inversions have a higher relative probability of fixation on Y chromosomes than on autosomes when *s* = 0.002 (*Ns* = 20; Figure S5) — in both cases, these average fixation probabilities stay lower than neutral. As these values of *Ns* often lie in the bulk of estimated DFEs, it appears difficult to draw firm conclusions on whether inversions should fix more easily on autosomes or on sex chromosomes. The relative fixation probabilities of Y-linked and autosomal inversions also depend on the distribution of dominance coefficients of deleterious mutations, on which little is known. Studies of spontaneous and induced mutations in *Drosophila* indicate that while lethal mutations are nearly (but not fully) recessive on average (*h* ≈ 0.02), mildly selected mutations have higher dominance coefficients: Simmons and Crow (1977) estimated that *h* ~ 0.3 on average for those mutations, while the more recent analysis of Manna et al. (2011) — that also included data from *S. cerevisiae* and *C. elegans* — estimated that *h* ~ 0.27 on average. Yeast gene knockout data have confirmed that mutations with a strong homozygous effect tend to have lower dominance coefficients than weak-effect mutations, the heterozygous effect being of the same order of magnitude for both types of mutations (Agrawal and Whitlock, 2011; Manna et al., 2012). These data were also used by Agrawal and Whitlock (2011) to infer the joint distribution of selection and dominance coefficients of mutations, indicating that while strongly deleterious mutations tend to be partially recessive, weakly deleterious mutations tend to have dominant effects. This last result should be taken with caution, however (as acknowledged by Agrawal and Whitlock), given the limited power associated with measures of fitness effects of weakly selected mutations. Overall, more data are thus needed to better estimate the proportion of mutations with very low dominance coefficients — of course, the dominance level of deleterious mutations on proto-Y chromosomes should be considered with caution, should these data become available, as the evolution of Y silencing would make mutations appear as more recessive then they actually are when recombination suppression first evolves (Lenormand et al., 2020). Nevertheless, these data and the results shown in the present article indicate that deleterious mutations are unlikely to generate a net fitness benefit for Y-linked inversions, as a high proportion of mutations should be very recessive for average fixation probabilities of such inversions to be greater than neutral. Our results also show that the net fitness benefit generated by very recessive mutations disappears above a threshold population size (Figure S12), and may be reduced by recurrent deleterious mutations occurring at the same locus (Figure S13). While mutation rates par nucleotide are sufficiently low so that the effect of recurrent mutations at the same nucleotide should be negligible, recurrent mutations within the same gene may have similar effects in the absence of complementation among these mutations (e.g., Johri and Charlesworth, 2025), potentially limiting the advantage of Y-linked inversions in the presence of very recessive mutations.

### Is the sheltering effect a potentially important component of sex chromosome evolution?

The sheltering of deleterious alleles facilitates the fixation of Y-linked inversions in regimes where the occurrence of a mutation-free inversion is extremely unlikely. However, and as mentioned above, sheltering is not the driver of the spread of inversions: in our model (and as in Olito et al., 2024; Jay et al., 2022, 2025), inversions tend to spread because they carry a lower-than-average mutation load, while sheltering reduces the disadvantage of initially carrying mutations. In addition, our results show that the sheltering effect concerns inversions that exceed a certain size (that depends on parameter values), while smaller inversions can fix only if they are initially free of deleterious mutation. The sheltering effect is thus not a necessary condition for the evolution of non-recombining sex chromosomes: if sheltering was not operating, recombination arrest would also occur through the spread of lucky inversions, but would involve smaller strata in regimes where a substantial proportion of deleterious mutations have weak fitness effects. Recombination arrest may also be caused by sex-antagonistic (SA) loci, which may greatly increase the fixation probability of X or Y-linked inversions even when their fitness effect is weak, and thus difficult to detect (Flintham and Mullon, 2025). Furthermore, the effect of deleterious mutations alone is not sufficient to explain the maintenance of evolutionary strata on sex chromosomes: indeed, recombination arrest causes mutation accumulation and a decline in fitness of the heterogametic sex, favoring the restoration of recombination in order to eliminate accumulated mutations (Lenormand and Roze, 2024). Another mechanism is thus needed to stably maintain recombination arrest, and avoid a strong drop in fitness of the heterogametic sex as the size of the non-recombining region increases. As shown by Lenormand and Roze (2022), the early evolution of Y gene silencing and dosage compensation can explain the stable maintenance of recombination arrest and the evolution of degenerate Y or W chromosomes. Sheltering is thus neither a necessary nor a sufficient component of sex chromosome evolution; nevertheless, it may play a significant role by helping larger inversions to reach fixation, thereby accelerating the process.

### Does inbreeding generate an advantage to inversions linked with a permanently heterozygous allele?

Our results show that selfing has two opposing effects on the spread of inversions capturing a mating-type locus. Both effects stem from the excess homozygosity caused by selfing, this excess being suppressed in an inversion linked with the mating-type locus. This enforced heterozygosity reduces the contribution of homozygous mutations to the mutation load, generating an advantage for such inversions (through the masking of deleterious mutations, corresponding to a sheltering effect), that becomes stronger when mutations are more recessive. However, decreasing homozygosity also reduces the efficiency of selection against deleterious alleles, so that at mutation-selection balance, the marginal fitness of the inversion is lower than the marginal fitness of an equivalent recombining segment. This generates a disadvantage for inversions capturing the mating-type locus, due to the less efficient purging of mutations. The importance of this purging disadvantage (relative to the masking advantage) increases with the strength of selection against deleterious alleles (as mutation-selection balance is reached more rapidly as *s* increases). As a consequence, we found that inversions capturing a mating-type locus can hardly fix under complete selfing and when *N*_e_ *s* is large (except when mutations are very recessive), while deleterious alleles accumulate in the population when *N*_e_ *s* is not large, due to the reduced effect of recombination caused by selfing. Given the selfing rates inferred in *Microbotryum* fungi reproducing by intratetrad mating (ranging from 0.5 to 0.9, Vercken et al., 2010; Abbate et al., 2018), our simulation results indicate that the sheltering of deleterious mutations may have helped the fixation of inversions extending the non-recombining regions associated with mating-type loci in these species (Badouin et al., 2015; Branco et al., 2018; Hartmann et al., 2021). In the case of XY or ZW sex chromosomes, inbreeding should have similar effects to those of selfing described here (generating both a masking advantage and a purging disadvantage for Y- or W-linked inversions) so that inbreeding may either increase or decrease the probability of fixation of inversions, depending on the distribution of selection and dominance of deleterious mutations. These effects would be worth exploring in greater detail.

## Conclusion

Overall, deleterious mutations play a key role in the evolution of sex chromosomes. They are at the basis of the process of degeneration and dosage compensation that are are probably necessary for the long term-maintenance of heteromorphic sex chromosomes. They also generate variance in fitness among copies of the same chromosome segment in different individuals, which can lead to the fixation of lucky inversions benefitting from a temporary fitness advantaged caused by a lower than average mutation load. The spread of inversions extending a permanently heterozygous sex-determining region is also affected by two antagonistic processes caused by deleterious mutations, whose relative importance depends on parameter values: their sheltering (notably when they have weak fitness effects and are very recessive), and their accumulation caused by the lack of recombination. In the long term, mutation accumulation should offset any initial advantage of recombination arrest, however, unless the effect of mutations can be silenced.

## Data availability

the source code of C++ simulation programs are available on Zenodo: https://doi.org/10.5281/zenodo.17287166.

## Acknowledgements

We thank Brian Charlesworth, Tatiana Giraud, Colin Olito, Amandine Véber and two anonymous reviewers for helpful discussions and comments, and the bioinformatics and computing services at Roscoff’s Biological Station (Abims platform) for computing time. This project has received funding from ERC RegEvol no. 101097167.

