## Supplementary Figures for "Effects of deleterious mutations on the fixation of chromosomal inversions on autosomes and sex chromosomes"

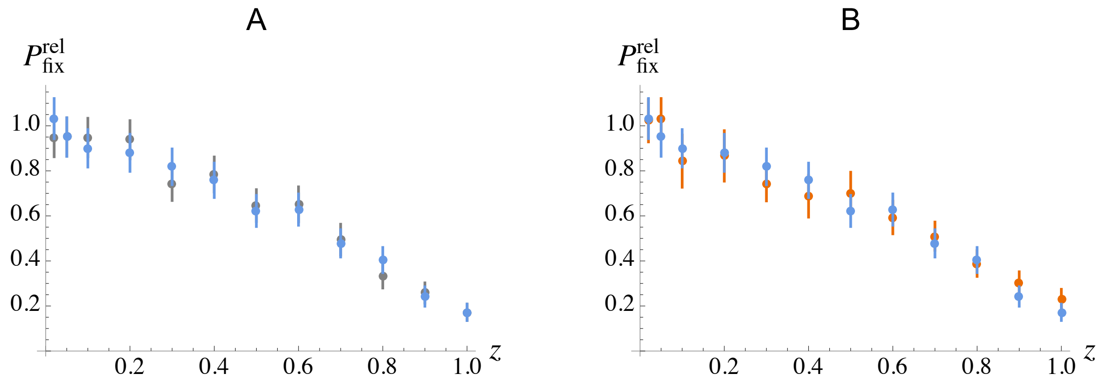


**Figure S1.** Blue dots in A and B show the same results as in Figure 1A (fixation probabilities of autosomal inversions relative to neutral, from multilocus simulations). Grey dots in A show fixation probabilities obtained in the absence of mutation segregating in the portion of the chromosome not encompassed by the inversion (in order to remove any possible linked selection effect), for the same parameter values. Red dots in B show fixation probabilities obtained when deleterious mutations occur at $L = 1{,}000$ discrete sites regularly spaced along the chromosome, with equal forward and backward mutation rate $u = U/L = 10^{-4}$ at each site.

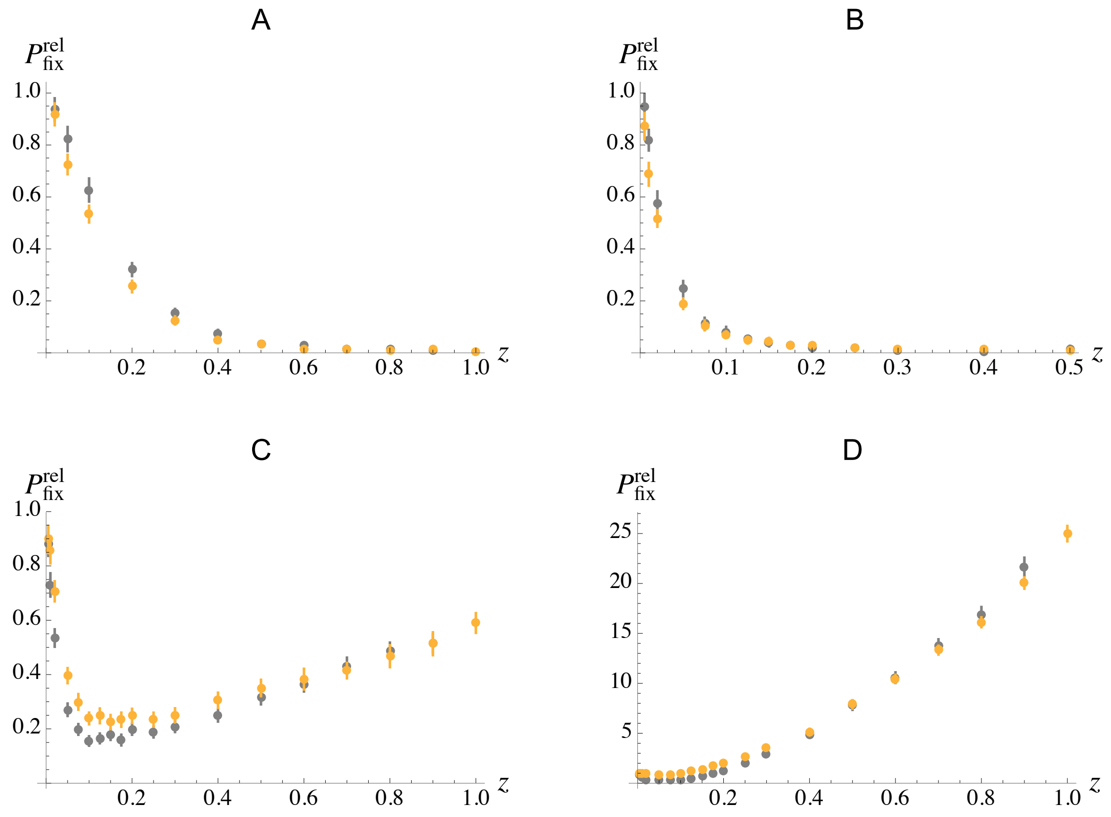


**Figure S2.** Mean fixation probabilities of Y-linked inversions (relative to neutral), as a function of their size $z$ relative to the total length of the chromosome. Orange: standard multilocus simulations; grey: fixation probabilities obtained in the absence of mutation segregating in the portion of the chromosome not encompassed by the inversion (in order to remove any possible linked selection effect). Parameter values are $N = 10^4$, $U = 0.1$, $R = 0.5$, $s = 0.05$, $h = 0.25$ (A), $s = 0.01$, $h = 0.25$ (B), $s = 0.125$, $h = 0.02$ (C), $s = 0.625$, $h = 0.004$ (D).

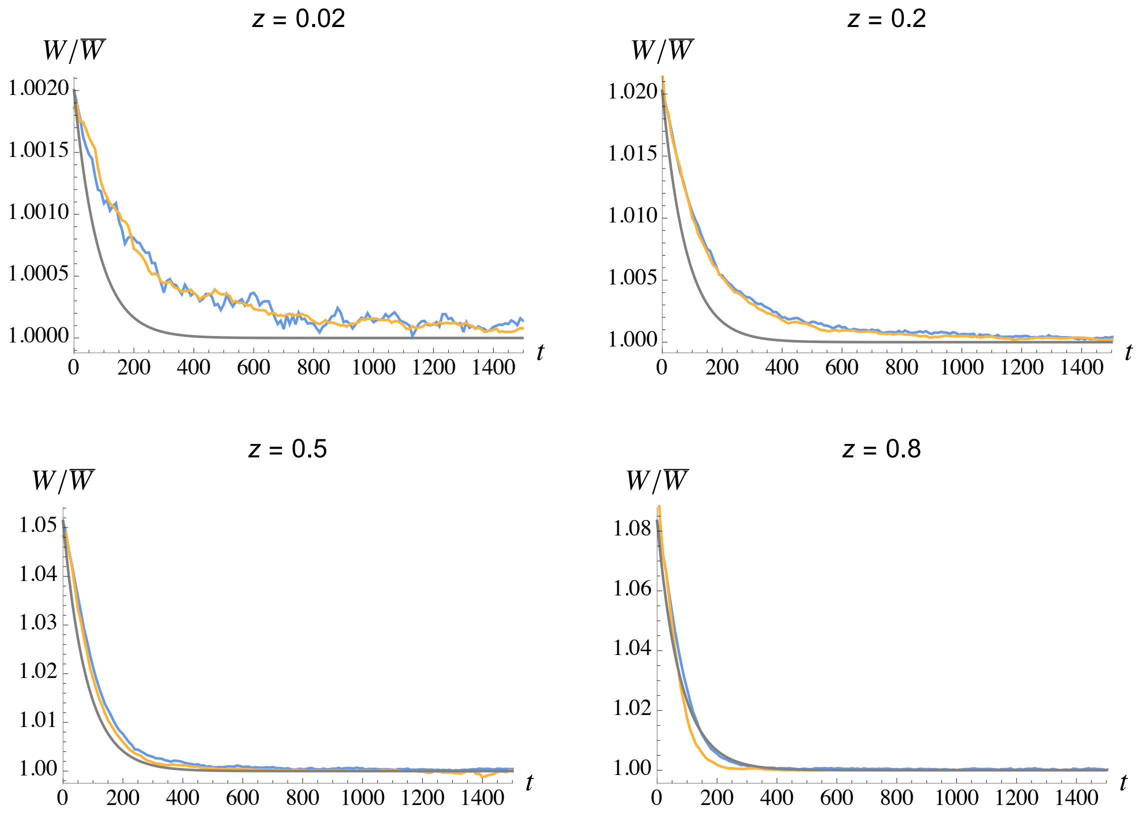


**Figure S3.** Average relative marginal fitnesses over time of inversions that reach fixation, for different inversion sizes $z$. The numbers of autosomal inversions (blue curves) that have reached fixation are 516, 440, 311 and 202 (over $10^7$ trials) for $z = 0.02$, 0.2, 0.5 and 0.8 (respectively). The number of Y-linked inversions (orange curves) that have reached fixation are 1836, 511, 64 and 18 for $z = 0.02$, 0.2, 0.5 and 0.8 (also over $10^7$ trials). As in Figure 1B, grey curves show the deterministic prediction $e^{Uz\,e^{-sht}}$ (from Nei et al., 1967).

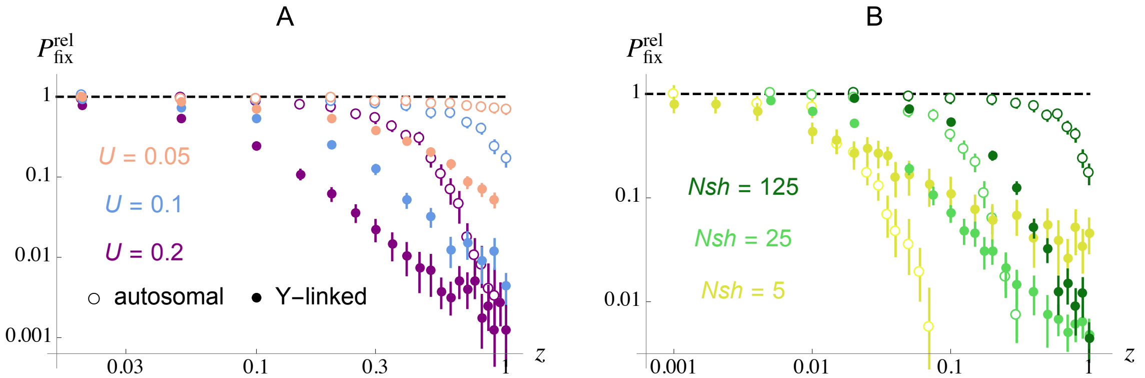


**Figure S4.** Same as Figure 4, where relative fixation probabilities are shown as a function of inversion size $z$ (relative to the size of the chromosome), on a log scale.

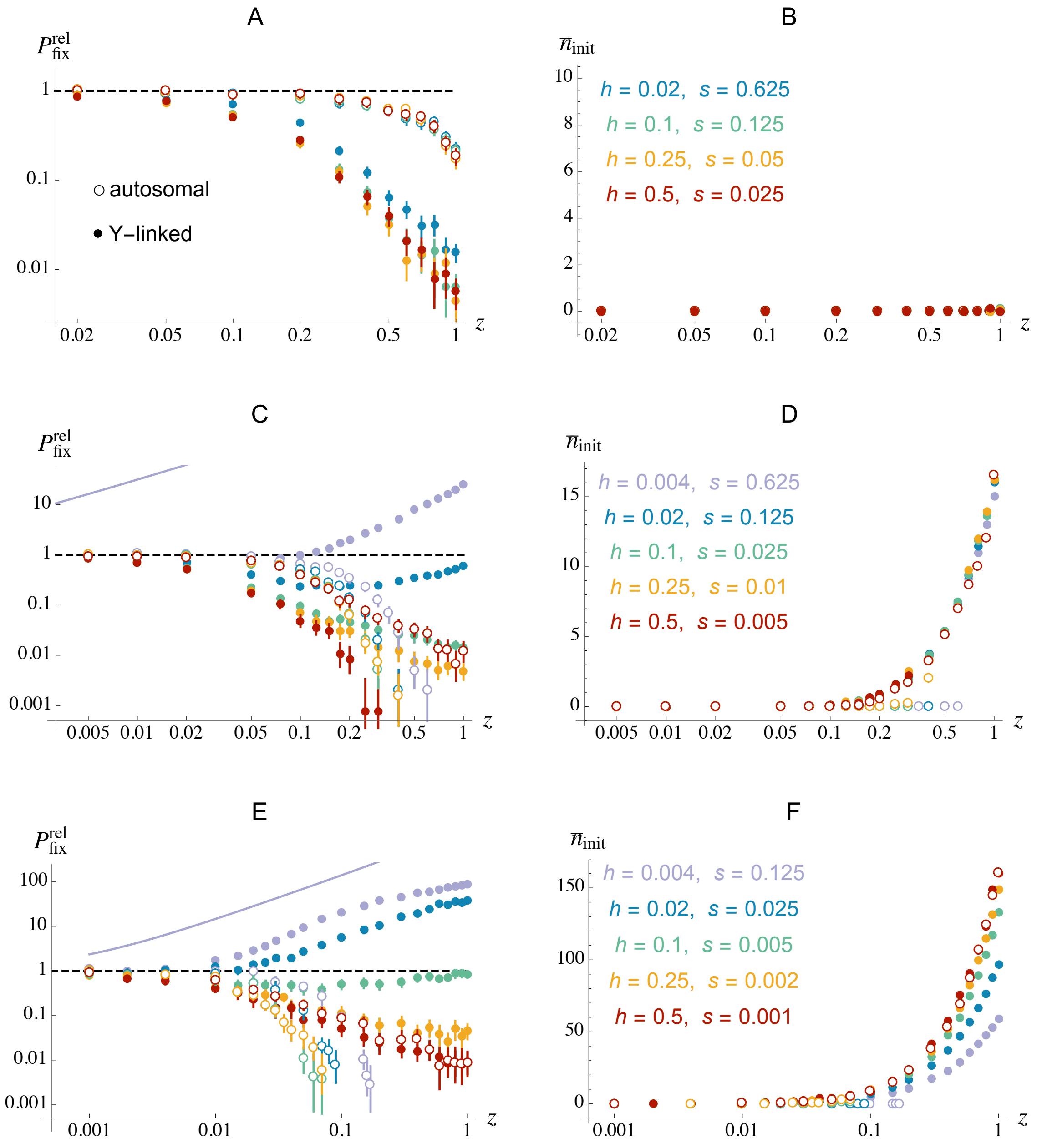


**Figure S5.** Same as Figure 5, where relative fixation probabilities (A, C, E) and average number of deleterious mutations initially present in inversions that reach fixation (B, D, F) are shown as a function of inversion size $z$ (relative to the size of the chromosome), on a log scale. Parameter values: $Nsh = 125$ (A, B), $Nsh = 25$ (C, D) and $Nsh = 5$ (E, F), $N = 10^4$, $U = 0.1$, $R = 0.5$. Curves in C, E show the upper bound for the fixation probability of Y-linked inversions provided by equation 2 of Charlesworth and Olito (2024) when $h = 0$ and $N_e s \gg 1$.

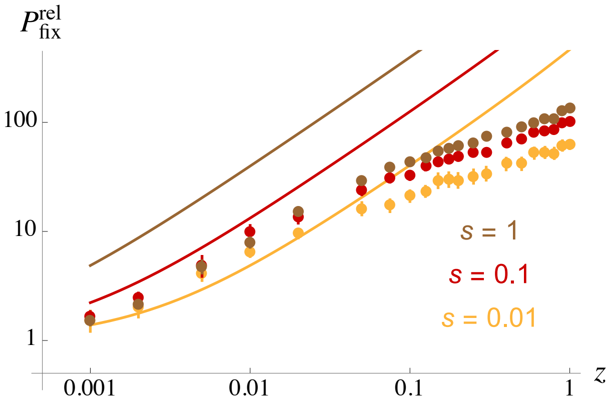


**Figure S6.** Relative fixation probabilities of Y-linked inversions as a function of their size $z$ (relative to the size of the chromosome), in the presence of fully recessive deleterious mutations ($h = 0$), for different values of $s$. Curves show the upper bound provided by equation 2 of Charlesworth and Olito (2024) when $h = 0$ and $N_e s \gg 1$. Other parameter values are as in Figure 5: $N = 10^4$, $U = 0.1$, $R = 0.5$.

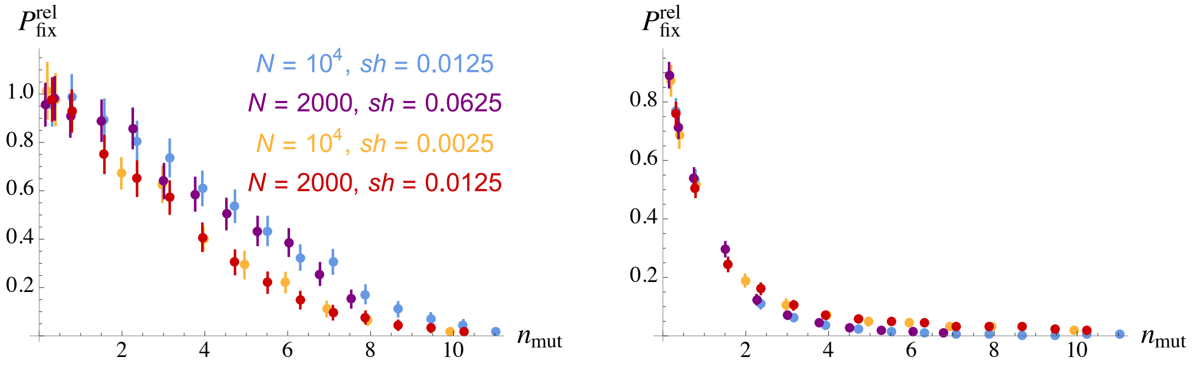


**Figure S7.** Mean fixation probabilities of autosomal (left) and Y-linked (right) inversions (relative to neutral), as a function of the average number of deleterious mutations in the population on a chromosome segment of the same size as the inversion ($n_{\rm mut}$), for different values of $N$ and $sh$. Note that the $Nsh$ product is the same for purple and blue dots ($Nsh = 125$) and for orange and red dots ($Nsh = 25$). Parameter values are $s = 0.05$, $h = 0.25$, $U = 0.2$, $R = 1$ (blue); $s = 0.125$, $h = 0.5$, $U = 0.5$, $R = 2.5$ (purple); $s = 0.01$, $h = 0.25$, $U = 0.1$, $R = 0.5$ (orange); $s = 0.05$, $h = 0.25$, $U = 0.2$, $R = 1$ (red).

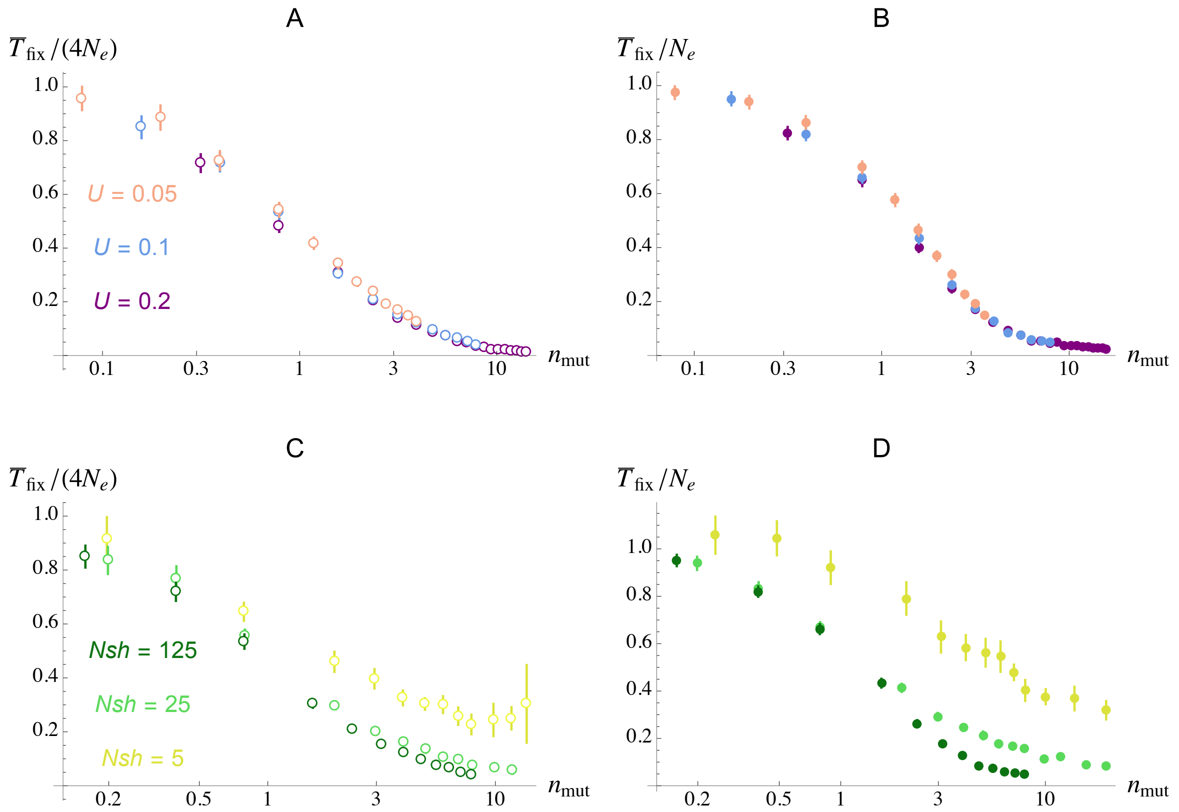


**Figure S8.** Mean fixation time of inversions as a function of the average number of deleterious mutations in the population on a chromosome segment of the same size as the inversion ($n_{mut}$), for the same parameter values as in Figure 4. A, C: autosomal inversions; B, D: Y-linked inversions. Mean fixation times are divided by the mean fixation time of a neutral mutation, $4N_e$ in the case of autosomal inversions and $N_e$ in the case of Y-linked inversions. For this, effective population size $N_e$ is computed as $N_e = N e^{-2U/R}$, using equation 9 in Hudson and Kaplan (1995) which approximates the effect of background selection when $sh \ll R$ and $N_e sh \gg 1$ (see Methods).

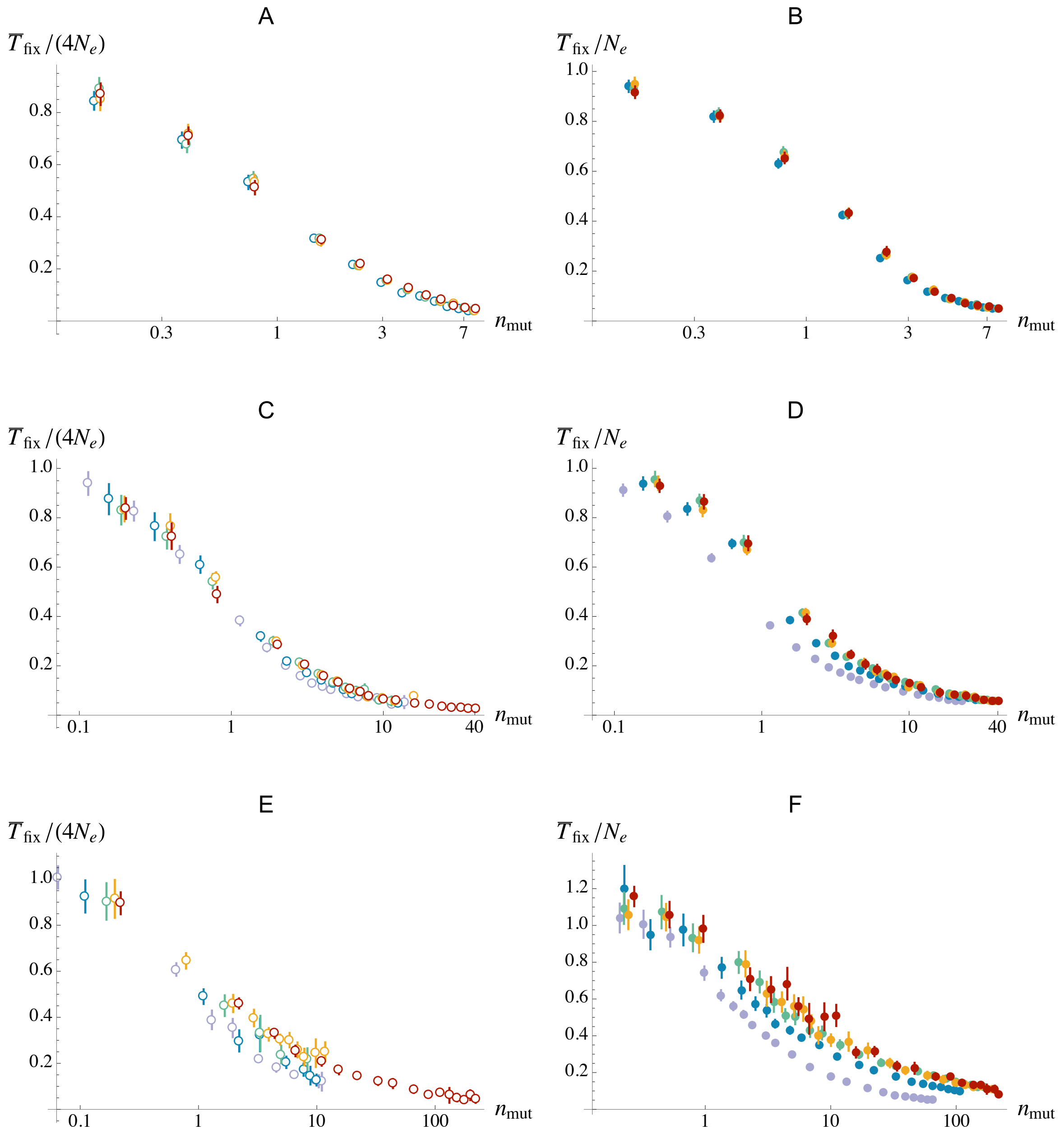


**Figure S9.** Mean fixation time of inversions as a function of the average number of deleterious mutations in the population on a chromosome segment of the same size as the inversion ($n_{\text{mut}}$), for the same parameter values as in Figure 5. A, C, E: autosomal inversions; B, D, F: Y-linked inversions. A, B: $Nsh = 125$; C, D: $Nsh = 25$; E, F: $Nsh = 5$. Mean fixation times are divided by the mean fixation time of a neutral mutation, $4N_{\text{e}}$ in the case of autosomal inversions and $N_{\text{e}}$ in the case of Y-linked inversions. For this, effective population size $N_{\text{e}}$ is computed as $N_{\text{e}} = N e^{-2U/R}$, using equation 9 in Hudson and Kaplan (1995) which approximates the effect of background selection when $sh \ll R$ and $N_{\text{e}} sh \gg 1$ (see Methods).

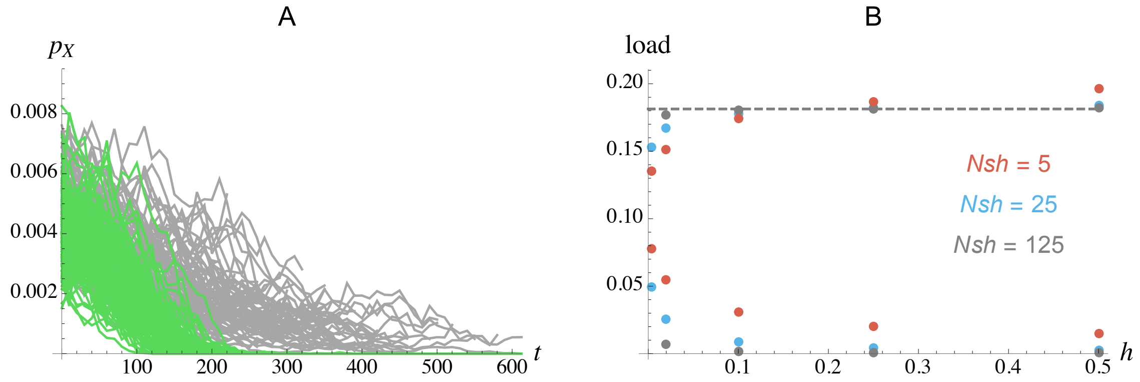


**Figure S10.** A: green: average frequency on X chromosomes over time of deleterious alleles initially present on Y chromosome inversions that reach fixation (each curve corresponds to the average over all mutations initially present in one successful inversion). Grey: same quantity measured for an equivalent segment of a randomly sampled Y chromosome (not carrying the inversion) at the time where the inversion first occurs. Parameter values: $N = 10^4$, $U = 0.1$, $R = 0.5$, $s = 0.625$, $h = 0.004$. Deleterious mutations initially present in a successful inversion are eliminated more rapidly from the population of X chromosomes than other mutations. B: top points: average mutation load measured in the multilocus simulations (in the case of a chromosome carrying the sex determining locus), in the absence of any inversion. The load is measured as $1 - \overline{W}$, where $\overline{W}$ is mean fitness. The different colors correspond to different values of $Nsh$. Bottom points: load generated by homozygous mutations (measured from the average reduction in fitness of individuals caused by homozygous mutations). Error bars are smaller than the size of symbols. Dashed line: deterministic approximation for the load when $h$ is not too small, $1 - e^{-2U}$. Parameter values are $N = 10^4$, $U = 0.1$, $R = 0.5$, $sh = 0.0005$ (red), $sh = 0.0025$ (blue), $sh = 0.0125$ (grey). Simulations have been performed for $h = 0.5$, 0.25, 0.1, 0.02 and 0.004 (last value only for $sh = 0.0025$ and $sh = 0.0005$). A substantial fraction of the load is caused by homozygous mutations for the lowest values of $h$ considered, particularly when $Nsh$ is small.

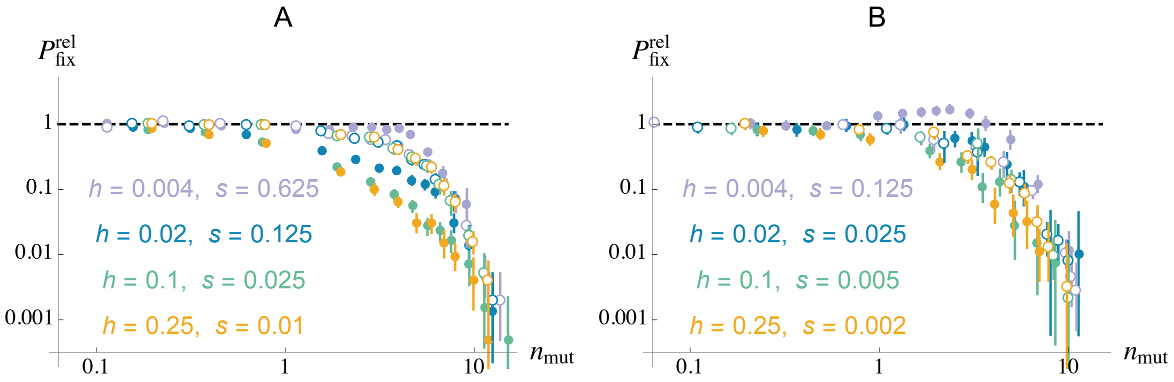


**Figure S11.** Equivalent of Figure 5C (A, $Nsh = 25$) and Figure 5E (B, $Nsh = 5$), where the fixation of inversions that initially carry deleterious mutations is not taken into account in the computation of fixation probabilities (results are shown only in the case of partially recessive mutations, $h < 0.5$). This shows that the relative fixation probability of Y-linked inversions that are initially mutation-free is generally lower or similar to the fixation probability of autosomal, mutation-free inversions (except for $h = 0.004$). The increased relative fixation probability of Y-linked inversions shown in Figures 5C and 5E (relative to autosomal inversions) when $h < 0.5$ is therefore mostly due to the fixation of Y-linked inversions that initially carry deleterious mutations.

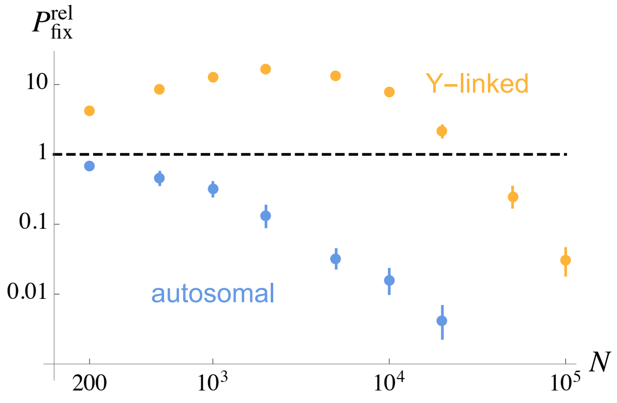


**Figure S12.** Relative fixation probabilities of autosomal (blue) and Y-linked (orange) inversions of size $z = 0.3$, as a function of population size $N$ (on a log scale), with $s = 0.625$, $h = 0.004$, $U = 0.1$ and $R = 0.5$.

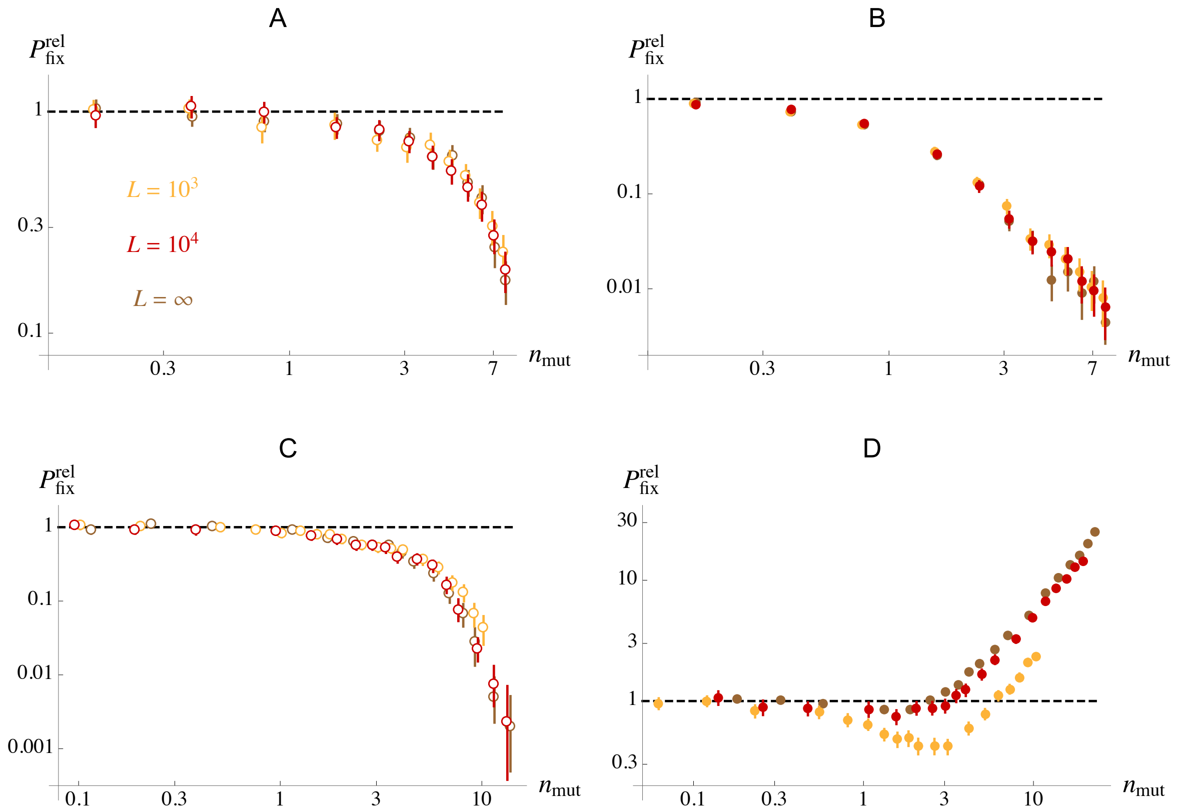


**Figure S13.** Relative fixation probabilities of autosomal (A, C) and Y-linked (B, D) inversions as a function of $n_{\rm mut}$, the average number of deleterious mutations in the population on a chromosome segment of the same size as the inversion (measured in the simulations), for $s = 0.05$, $h = 0.25$ (A, B) and $s = 0.625$, $h = 0.004$ (C,D). Brown dots have been obtained using our standard simulation program in which deleterious mutations occur at an infinite number of loci along the chromosome, while orange and red dots correspond to the results obtained when mutations occur at a finite number ($L$ = 1,000 and $L$ = 10,000, respectively) of loci regularly spaced along the chromosome, with equal forward and backward mutation rate $u = U/L$. Other parameter values are as in Figure 5: $N = 10^4$, $U = 0.1$, $R = 0.5$.

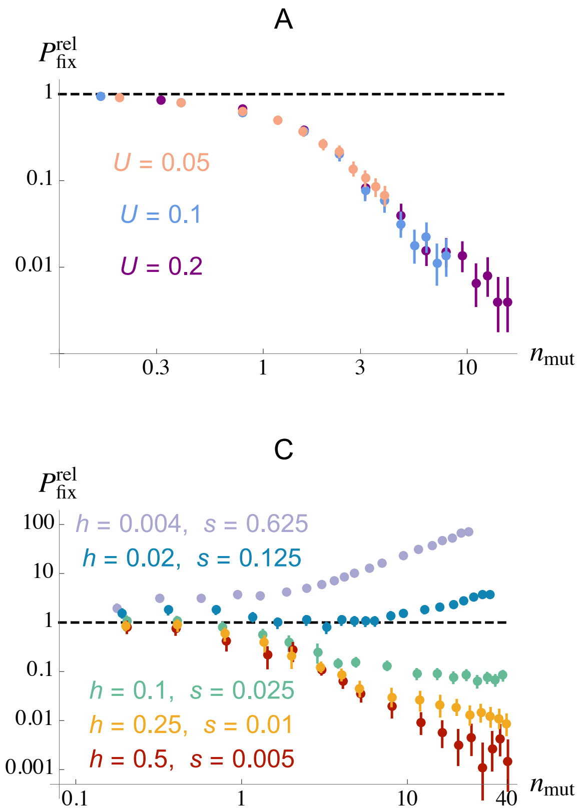


**Figure S14.** Relative fixation probabilities of inversions capturing a mating-type locus (in the absence of inbreeding), as a function of $n_{\text{mut}} \approx Uz/(sh)$, the average number of deleterious mutations in the population on a chromosome segment of the same size as the inversion, for different values of the deleterious mutation rate $U$ (A) and of the selection and dominance coefficients of deleterious alleles (B, C). $Nsh = 125$ in A, B, while $Nsh = 25$ in C. Parameter values are $s = 0.05$, $h = 0.25$ (in A), $U = 0.1$ (in B, C), $N = 10^4$. Chromosome map length $R$ is set to 0.5 when $U = 0.1$, and 0.25, 1 when $U = 0.05$, 0.2 (respectively) in order to maintain a constant $U/R$ ratio and thus a constant $N_e$ in the absence of segregating inversion (see Methods).

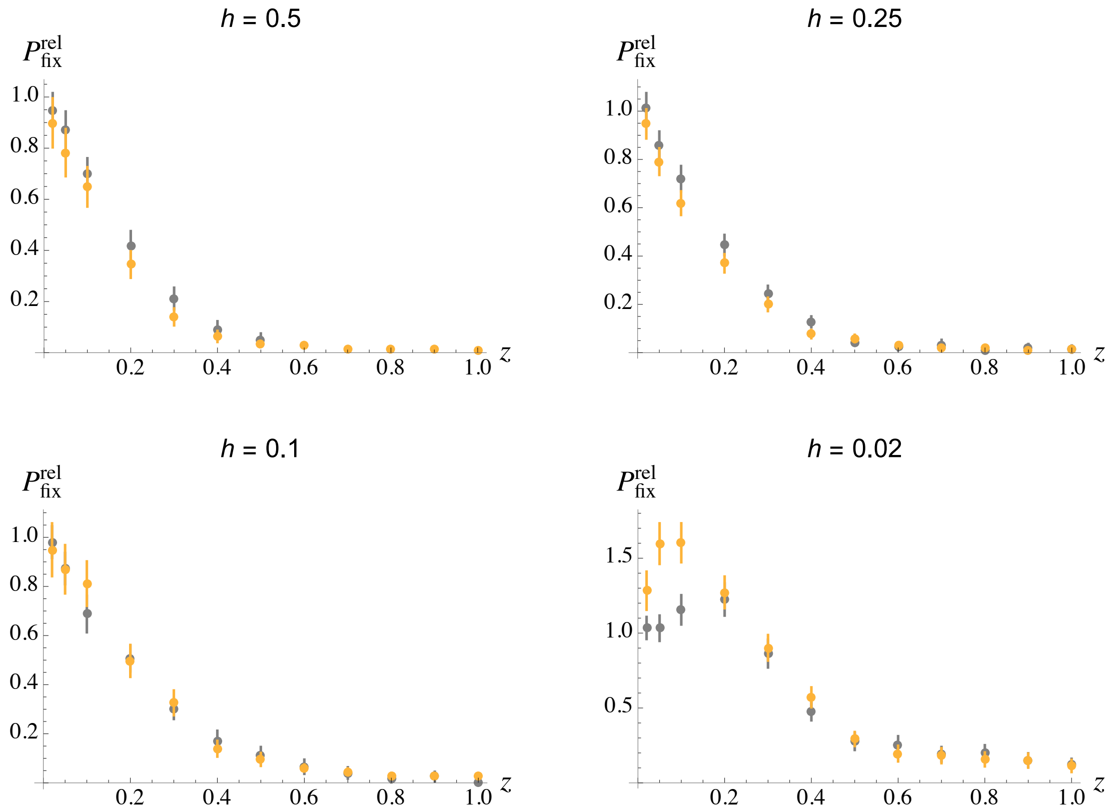


**Figure S15.** Relative fixation probabilities of inversions capturing a mating-type locus (in the absence of inbreeding), as a function of their size $z$ relative to the total length of the chromosome. Orange: standard multilocus simulations; grey: fixation probabilities obtained in the absence of mutation segregating in the portion of the chromosome not encompassed by the inversion (in order to remove any possible linked selection effect). Parameter values are $N = 10^4$, $U = 0.1$, $R = 0.5$, $Nsh = 125$.

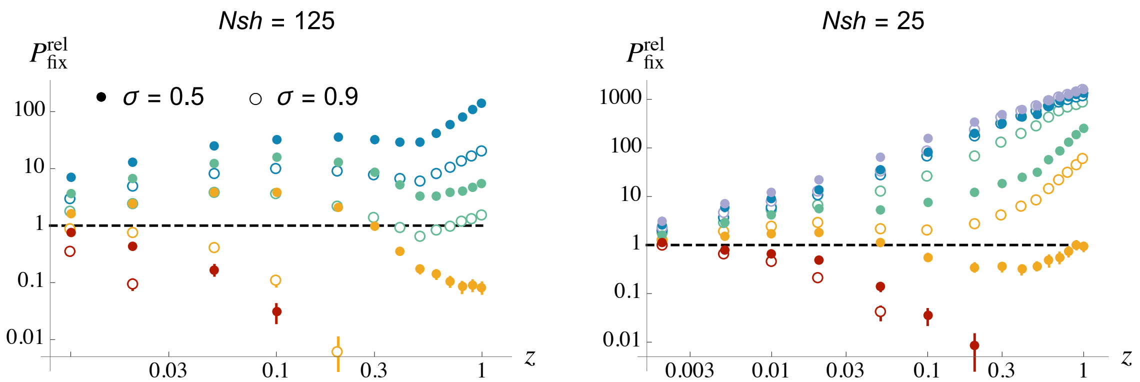


**Figure S16.** Relative fixation probabilities of inversions capturing a mating-type locus as a function of their size $z$, for different values of the selfing rate $\sigma$, and different selection and dominance coefficients of deleterious alleles. Same results as on Figure 7 (under regular selfing only), with $\sigma = 0.5$ and $\sigma = 0.9$ shown on the same figures.

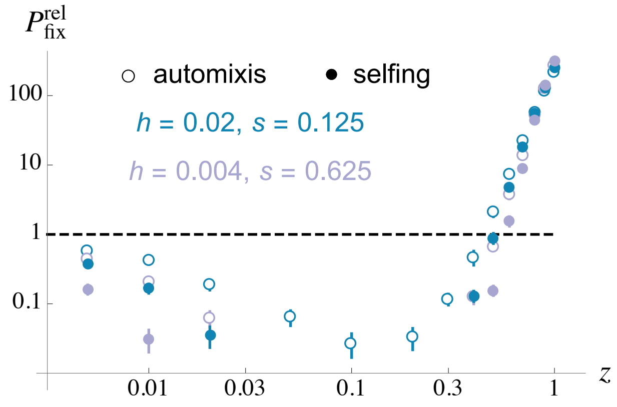


**Figure S17.** Relative fixation probabilities of inversions capturing a mating-type locus as a function of their size $z$, under complete selfing or automixis ($\sigma = 1$). For $Nsh = 25$, mutations accumulated rapidly in the population (in the absence of inversion) for $h = 0.5$, 0.25 and 0.1, and we could thus obtain results only for $h = 0.02$ and $h = 0.004$ (shown here). For $Nsh = 125$, mutations accumulated only for $h = 0.5$; however, for the other values of $h$ (0.25, 0.1, 0.02), no fixation of inversion occurred over the $10^7$ trials, except for $h = 0.25$ and the smallest inversion sizes ($z \leq 0.05$, not shown).

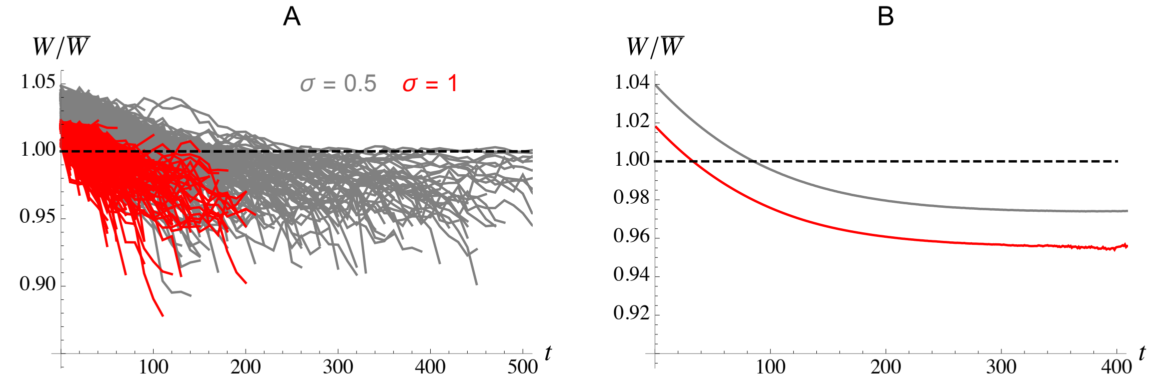


**Figure S18.** A. Relative marginal fitness over time of 500 inversions of size $z = 0.5$ capturing a mating-type locus, initially mutation-free, under intermediate selfing ($\sigma = 0.5$, grey) and complete selfing ($\sigma = 1$, red). Parameter values are $s = 0.05$, $h = 0.25$, $N = 10^4$, $U = 0.1$, $R = 0.5$. For these parameter values, mutations do not accumulate in the population even when selfing is complete. B. mean relative marginal fitness over time of inversions of size $z = 0.5$ capturing a mating-type locus, initially mutation-free, for the same parameter values. Averages are taken over $\sim 10^7$ trajectories for $\sigma = 0.5$ and over $\sim 10^9$ trajectories for $\sigma = 1$, the average at time $t$ being measured over inversions that are still segregating at time $t$ ($\sim$ 267,000 inversions were still segregating at generation 400 for $\sigma = 0.5$, but only 1,070 for $\sigma = 1$).

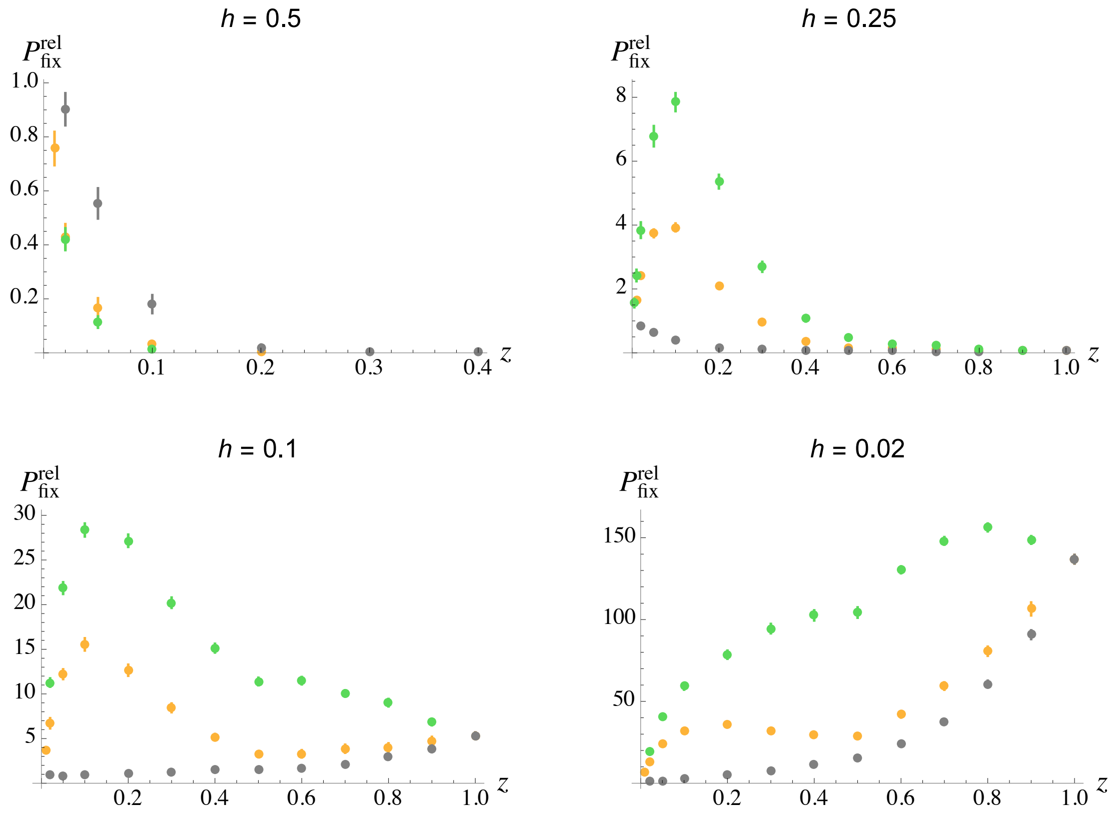


**Figure S19.** Relative fixation probabilities of inversions capturing a mating-type locus as a function of their size $z$, with a selfing rate $\sigma = 0.5$. Orange: standard multilocus simulations; grey: fixation probabilities obtained in the absence of mutation segregating in the portion of the chromosome not encompassed by the inversion (in order to remove any possible linked selection effect); green: fixation probabilities obtained when the map length outside the inversion is set to $R(1-z)$ (instead of $R$) in heterozygous individuals for the inversion (i.e., the inversion does not increase recombination outside the inversion in heterokaryotypes). Parameter values are $N = 10^4$, $U = 0.1$, $R = 0.5$, $Nsh = 125$.

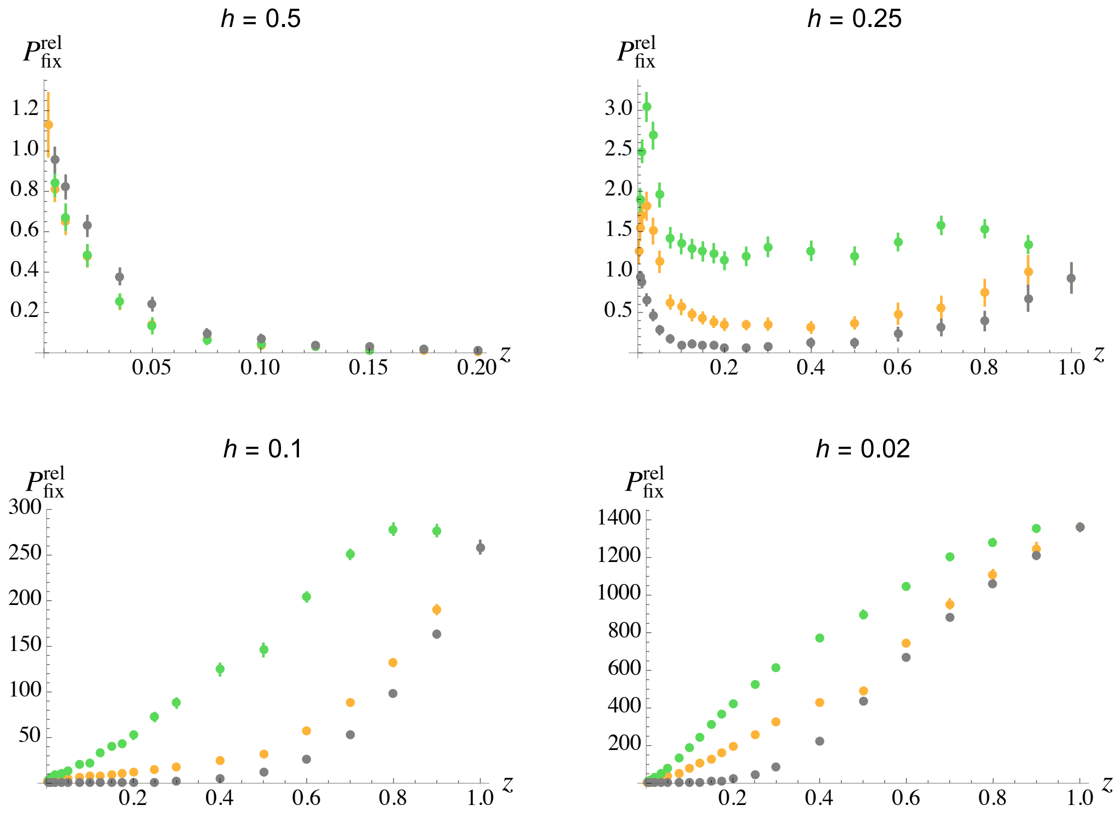


**Figure S20.** Same as Figure S19 with $Nsh = 25$.
